# Admixture/fine-mapping in Brazilians reveals a West African associated potential regulatory variant (rs114066381) with a strong female-specific effect on body mass- and fat mass-indexes

**DOI:** 10.1101/827311

**Authors:** Marilia O Scliar, Hanaisa P Sant Anna, Meddly L Santolalla, Thiago P Leal, Nathalia M Araújo, Isabela Alvim, Victor Borda, Wagner CS Magalhães, Mateus H Gouveia, Ricardo Lyra, Moara Machado, Lucas Michelin, Maíra R Rodrigues, Gilderlanio S Araújo, Fernanda SG Kehdy, Camila Zolini, Sérgio Viana Peixoto, Marcelo Luizon, Francisco Lobo, Michel S Naslavsky, Guilherme L Yamamoto, Yeda A O Duarte, Matthew EB Hansen, Shane A Norris, Robert H Gilman, Heinner Guio, Ann W Hsing, Sam M Mbulaiteye, James Mensah, Julie Dutil, Meredith Yeager, Edward Yeboah, Sarah A Tishkoff, Ananyo Choudhury, Michele Ramsay, Maria Rita Passos-Bueno, Mayana Zatz, Timothy D O’Connor, Alexandre C Pereira, Mauricio L Barreto, Maria Fernanda Lima-Costa, Bernardo L Horta, Eduardo Tarazona-Santos

## Abstract

Admixed populations are a resource to study the global genetic architecture of complex phenotypes, which is critical, considering that non-European populations are severely under-represented in genomic studies. Leveraging admixture in Brazilians, whose chromosomes are mosaics of fragments of Native American, European and African origins, we used genome-wide data to perform admixture mapping/fine-mapping of Body Mass Index (BMI) in three population-based cohorts from Northeast (Salvador), Southeast (Bambuí) and South (Pelotas) of the country. We found significant associations with African-associated alleles in children from Salvador (PALD1 and ZMIZ1 genes), and in young adults from Pelotas (NOD2 and MTUS2 genes). More importantly, in Pelotas, rs114066381, mapped in a potential regulatory region, is significantly associated only in females (p= 2.76 e-06). This variant is very rare in Europeans but with frequencies of ~3% in West Africa, and has a strong female-specific effect (95%CI: 2.32-5.65 kg/m2 per each A allele). We confirmed this sex-specific association and replicated its strong effect for an adjusted fat-mass index in the same Pelotas cohort, and for BMI in another Brazilian cohort from São Paulo (Southeast Brazil). A meta-analysis confirmed the significant association. Remarkably, we observed that while the frequency of rs114066381-A allele ranges from 0.8 to 2.1% in the studied populations, it attains ~9% among morbidly obese women from Pelotas, São Paulo, and Bambuí. The effect size of rs114066381 is at least five-times the effect size of the FTO SNPs rs9939609 and rs1558902, already emblematic for their high effects, and for which we replicated associations in Pelotas. We demonstrate how, after a decade of GWAS mostly performed in European-ancestry populations, non-European and admixed populations are a source of new relevant phenotype-associated genetic variants.

## INTRODUCTION

Overweight and obesity are major risk factors for noncommunicable diseases, which are responsible for 63% of deaths in the world (WHO 2011) and for 72% in Brazil (Schmidt et al. 2011). Inter-individual differences in BMI result from the effects of multiple genetic variants, environmental factors, and their interactions (Locke et al. 2015; Rask-Andersen et al. 2017). Most of BMI heritability, estimated to be ~40%, is attributable to unknown genetic factors (Robinson et al. 2017, Wainschtein et al. 2019). For instance, a meta-analysis of genome-wide association studies (GWAS) of BMI estimated that 97 loci explain around 2.7% of its variance (Locke et al. 2015). The GWAS-Catalog (Welter et al. 2014) and the DANCE web tool (Araújo et al. 2016) report 389 SNPs associated with BMI in different populations, with a mean effect size of 0.054 kg/m^2^ (Figure S1). Thus, BMI genetic architecture is characterized by a high number of loci with small effect sizes (Robinson et al. 2017).

Our knowledge on the genetic architecture of complex phenotypes is biased (Sirugo et al. 2019) because only 22% of individuals included in GWAS are non-Europeans/non-US whites, while only 2.4% are from Africa and 1.3% from Latin America (Morales et al. 2018). For BMI, the meta-analysis by Locke et al. (2015) included only 5% of individuals of non-European ancestry among 339,226 individuals. Thus, expanding GWAS-based strategies beyond non-European populations is critical to discover differences in the genetic architecture of BMI among diverse populations. This is especially important for phenotypes such as obesity, whose prevalence is higher in US African Americans, Hispanics and Native Americans than in European Americans (Cheng et al. 2009; Nassir et al. 2012), and in Brazil, higher in black women than in white women (Gigante et al. 2009).

Another setback is that few studies consider the influences of age- and sex-associated genetic factors on BMI. Although there is a high correlation of intra-individual measurements of BMI at diverse ages, some genetic variants have distinct effects depending on age (Bradfield et al. 2012; Hardy et al. 2010; Warrington et al. 2015). For example, a meta-analysis of 14 GWAS (Graff et al. 2013) found that variants near to *PRKD1, TNNI3K, SEC16B*, and *CADM2* genes had larger effects on BMI during adolescence/young adulthood than later in the lifespan, while a variant near *SH2B1* had the opposite trend. The meta-GWAS by Winkler et al. (2015) of 114 studies with individuals of European descent identified 15 loci with different effects in individuals over- and under-50 years. Regarding sex, while Winkler et al. (2015) did not identify sex-dependent SNPs that affect BMI, Locke et al. (2015) found variants in *SEC16B* and *ZFP64* with stronger effects in women.

Here we study the genetic architecture of BMI in the admixed population of Brazil, the largest and most populous Latin-American country, with more than 200 million inhabitants. Brazilians are the product of about 500 years of admixture between Africans, Europeans and Native Americans (Kehdy et al. 2015) and therefore, are suitable for admixture mapping. This method uses an admixed population to map genomic regions associated both with a specific ancestry and the phenotype of interest. Admixture mapping, by performing less statistical tests respect to classical GWAS, results in higher statistical power. Thus, for medium-sized studies (more feasible in limited-resources setting environments hosting non-European populations), admixture mapping improves the power to detect an association when compared to GWAS that include only a few thousands of individuals. Admixture mapping may be followed by fine-mapping if high density data are available. So far, admixture mapping has identified seven loci associated with BMI at chromosomes 2, 3, 5, 15 and X (Basu et al. 2009; Cheng et al. 2009; Cheng et al. 2010), but these studies were restricted to US African American populations. In the admixed Brazilian population, admixture mapping may allow to find susceptibility variants associated with African or Native American ancestry, two groups still under-represented in genomic studies.

We performed admixture mapping (followed by fine-mapping) of BMI using data of ~2.5 million SNPs for three Brazilian population-based cohorts, from Northeast (Salvador), Southeast (Bambuí) and South (Pelotas) of Brazil, with distinct admixture and socio-demographic backgrounds, studied by the EPIGEN-Brazil Initiative (https://epigen.grude.ufmg.br/, Kehdy et al. 2015). Salvador has the largest African ancestry (51%) among the EPIGEN populations, and 43% and 6% of European and Native American ancestries, respectively. Differently, Pelotas and Bambuí have predominant European ancestry (76% and 79%, with 16%and 14% of African ancestry, and 8% and 7% of Native American ancestry, respectively) (Table S1, Figure 1). As these cohorts include individuals of three different epochs of life – children, young adults, and older adults – we performed three separate admixture mapping analyses to identify variants associated with BMI in each age group/cohort. Additionally, we performed a replication study by testing the association between 216 BMI GWAS-Catalog-hits in our three Brazilian cohorts.

**Figure 1.**
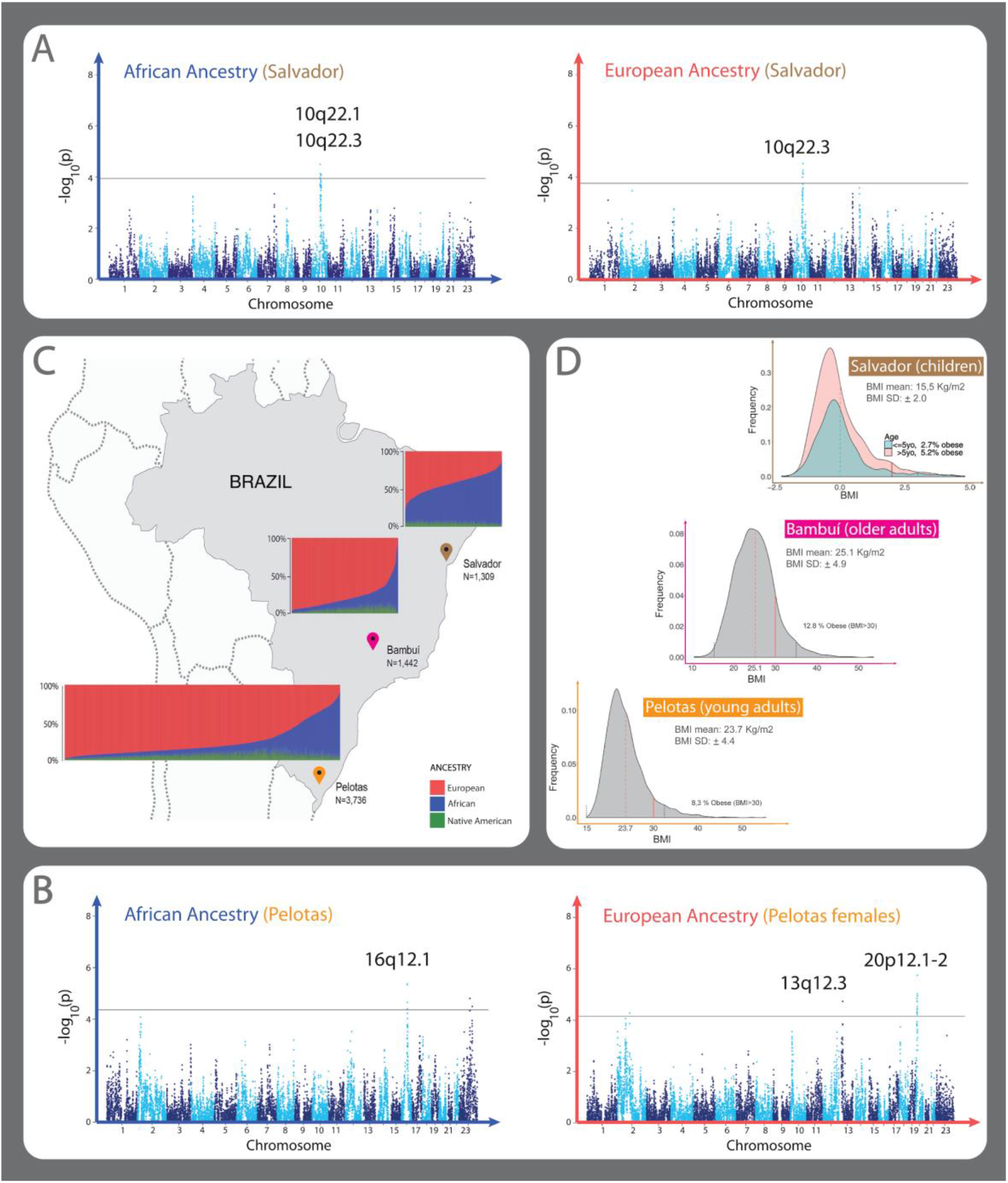
Admixture in the Brazilian cohorts, BMI distributions and Admixture Mapping (AM) Manhattan plots with significant peaks. **A-B** Manhattan plots showing AM peaks using linear regressions with PCAdmix local ancestry inferences. Consensus significant AM peaks for PCAdmix and RFMix local ancestry inferences are specified on each plot. **A.** Manhattan Plot showing the AM results of African (left) and European (right) ancestry in Salvador cohort. African ancestry AM shows two positive significant peaks 10q22.1 (β=0.36, p-value=3.21e-05) and 10q22.3 (β=0.36, p-value=7.87e-05). European AM ancestry analysis shows one negative associated peak 10q22.3 (β=-0.36, p-value=2.92e-05). **B.** Manhattan plot showing the AM results of African (left) and European (right) ancestry in the Pelotas cohort. One peak in 16q12.1 (β=-0.80, p-value=4.30e-06) was observed associated with African ancestry, and two associated peaks, 13q12.3 (β=-0.95, p-value=1.84e-05) and 20p12.1-2 (β=-1.05, p-value=1.79-06), with European ancestry in females. Results are presented as log_10_(p-value) to the given ancestry of each window of 100 SNPs along the genome. Black line in the Manhattan plots correspond to the genome-wide threshold p-value estimated for the given ancestry and dataset (Table S5). The linear regression coefficient (β) and p-values for all peaks correspond to the lead window, the genomic window with the most significant p-value in the linear regression result. **C**. Brazilian regions and continental individual ancestry bar plots for each cohort. **D.** Histogram of Z-score adjusted by sex and age according to WHO guidelines in Salvador (top), histogram of BMI in Bambui (center) and Pelotas (bottom) cohorts.

## RESULTS

### Admixture mapping and fine-mapping

We performed an admixture mapping analysis for the three continental ancestries (African, European, and Native American) in the three cohorts, adjusting for age (Salvador and Bambuí), sex, socioeconomic variables, and genome-wide African ancestry. Because in the Pelotas cohort we found an interaction between individual ancestry and sex (Tables S2, S3, S4), we also conducted sex-specific admixture mapping for this cohort. The Bambuí cohort includes a large number of relatives (Kehdy et al. 2015), thus we corrected for family structure (see Materials and Methods). We used an additive model considering the number of inferred African, European or Native American ancestry copies (0, 1 or 2) carried by an individual for each chromosome fragment.

Table 1 shows the five consensus significant admixture mapping peaks found in Salvador and Pelotas (i.e. p-value peaks for the same genomic regions using both PCAdmix (Brisbin et al. 2012) and RFmix (Maples et al. 2013) local ancestry inferences and using threshold p-values of Table S5). The distribution of BMI for each allele of African or European ancestry for the five peaks are shown in Supplementary Material (Figures S2, S3, S4). No consensus significant peak was found in older adults from Bambuí. We performed fine-mapping of these peaks using both genotyped and imputed data. For the fine-mapping, we considered as significant p-values less than or equal to the ones obtained for the admixture mapping peaks (Table 1), and suggestively significant (Tables S6, S7, Figure S5) those SNPs with a p-value higher than the ones obtained for the admixture mapping peaks but not more than one unit of -log (p-value).

**Table 1.** Admixture mapping peaks obtained both with RFMix and PCAdmix local ancestry inferences.

| Cohort | Genomic region<br>(length in base<br>pairs) | Local Ancestry<br>(Sub-dataset) | Regression<br>Coefficient<sup>a</sup> | P-value<sup>a</sup> | Associated Region<br>(length in base<br>pairs)<sup>b</sup> |
| --- | --- | --- | --- | --- | --- |
| CHILDREN<br>FROM<br>SALVADOR | 10q22.1<br>(4,300,000) | African | 0.36 | 3.21e-05 | 445,166 |
|  | 10q22.3<br>(4,400,000) | African | 0.36 | 7.87e-05 | 255,990 |
|  |  | European | -0.36 | 2.92e-05 | 577,546 |
| YOUNG<br>ADULTS<br>FROM<br>PELOTAS | 16q12.1<br>(5,600,000) | African | -0.80 | 4.30e-06 | 600,961 |
|  | 13q12.3<br>(3,300,000) | European<br>(female) | -0.95 | 1.84e-05 | 118,909 |
|  | 20p12.1-2<br>(8,700,000) | European<br>(female) | -1.05 | 1.79e-06 | 3,481,320 |

Regression coefficients, p-values, and windows length associated in the linear regressions using PCAdmix local ancestry inferences. <sup>a</sup>Regression coefficients and p-values of the lead genomic window of 100 SNPs, using PCAdmix inference. The lead window is the genomic window with the most significant regression coefficient in the Admixture Mapping among those windows below the significance cut-off. Regression coefficient are the change in units of BMI (Kg/m<sup>2</sup>) for each additional copy of a specific ancestry. <sup>b</sup>Length of the continuous genomic region including significant Admixture Mapping results. Each window of PCAdmix inference has 100 SNPs.

#### Fine-mapping on Salvador Children

The high African ancestry (51%) in children from Salvador allowed us to identify two genomic regions where this ancestry is positively associated with BMI and within these regions, we identified three significant SNPs (Tables S8, S9, Figure S5): within 10q22.1, rs1334909357-CTTT in an intron in the *PALD1* gene and, within 10q22.3, the linked SNPs rs79947827-A and rs141274185-T (linkage disequilibrium [LD]: r^2^= 0.86) in the *ZMIZ1* gene, that encodes a protein that regulates the activity of many transcription factors (Rogers et al. 2013). Other SNPs in *ZMIZ1* are associated with 19 complex disorders, and this gene is among the 21 human genes most associated with complex phenotypes, including not only BMI-related phenotypes such as height and sitting height ratio, but also psychiatric disorders, breast cancer and autoimmune diseases (http://gilderlanio.pythonanywhere.com/home, Figure S6).

#### Fine-mapping in Young adults from Pelotas

While the low non-European admixture reduces the power to detect non-European associated variants in Pelotas, this is compensated by its larger size (n=3,628) in respect to the Salvador cohort. Also, as Pelotas is a birth-cohort, all individuals have the same age, which limits non-genetic variance for BMI. For the entire Pelotas cohort, we identified one genomic region, 16q12.1, where African ancestry is negatively associated with BMI and, within this region, we found one significant SNP, rs76416629-G, 2kb upstream of *NOD2* gene (Tables S6, S7, Figure S5).

Furthermore, we identified a genomic region, 13q12.3, for which European ancestry is associated with lower values of BMI in females and, within this region, two significant SNPs (not in LD with each other, r^2^<0.001). rs113214936-G in the intron of *MTUS2* gene was associated with obesity-related traits by Comuzzie et al. (2012). Our most striking result is the significant association of the SNP rs114066381-A with a strong effect on BMI in females (beta=3.99±0.84 kg/m^2^ per allele, 95%CI: 2.32-5.65, p=2.76×10-6, Tables 2, Figure 2). This SNP is present in 31 unrelated females (all heterozygous) that have a mean BMI of 27.99 Kg/m^2^, which is larger than the mean BMI for the cohort females (23.61 Kg/m^2^, p=0.0008, Figure 3). These 31 females have a mean African ancestry of 35%, while the mean in unrelated females is 16%. Remarkably, the BMI of 25 males carrying the rs114066381-A allele (mean: 23.72 Kg/m^2^) does not differ from the general population (mean: 23.81 Kg/m^2^, p=0.5397, Figure 3).

**Figure 2.**
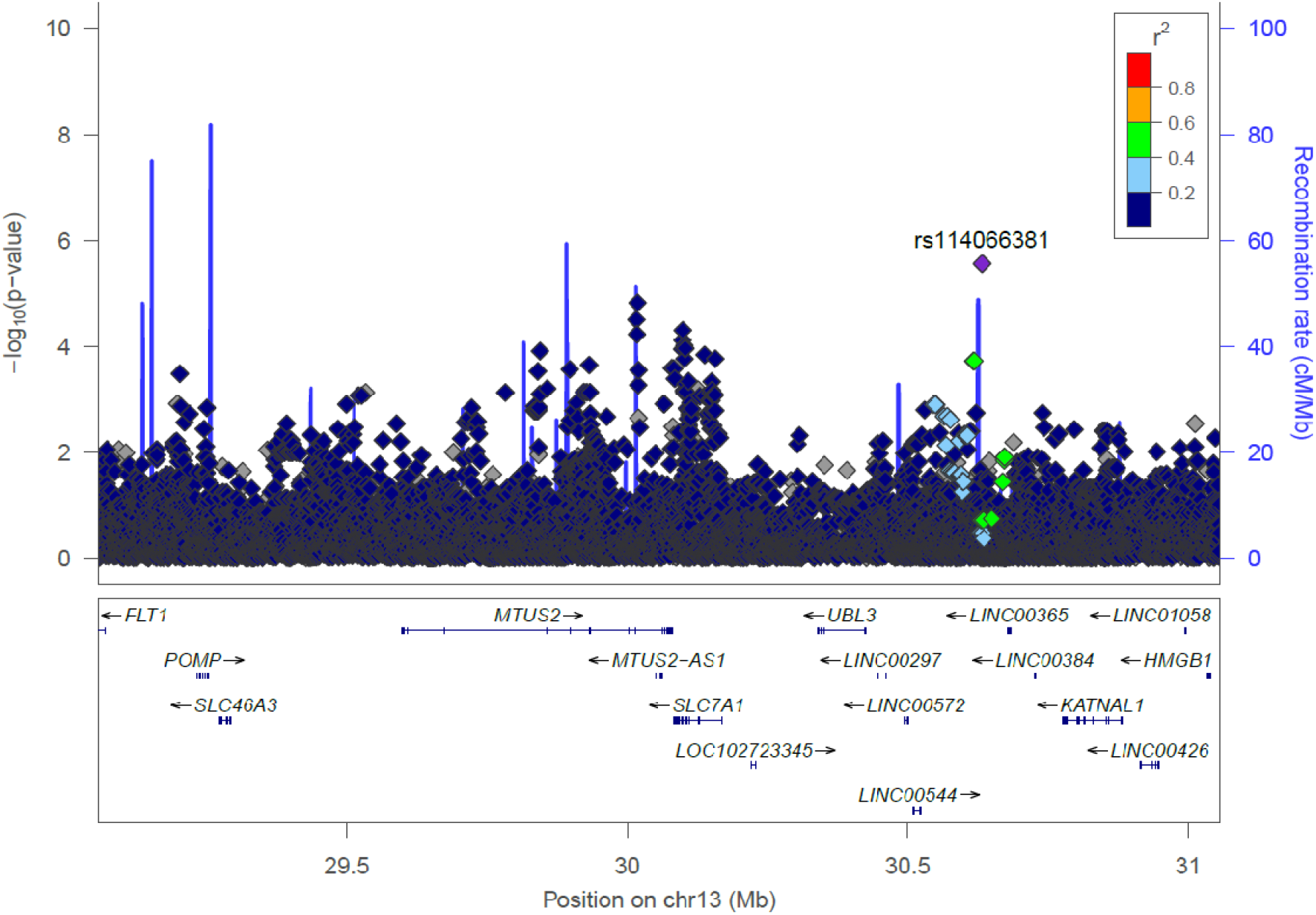
LocusZoom plot of the fine mapping of consensus significant Admixture Mapping peak in young adults from Pelotas at 13q12.3 associated with European ancestry in females performed using both genotyped and imputed SNPs ±1 Mb from target region (lead windows). The SNP with the lowest p-value is color coded in purple and labeled. The linkage disequilibrium between this SNP and the remaining nearby SNPs is indicated by the color coding according to r^2^ values based on Africans from 1000 Genomes Project.

**Figure 3.**
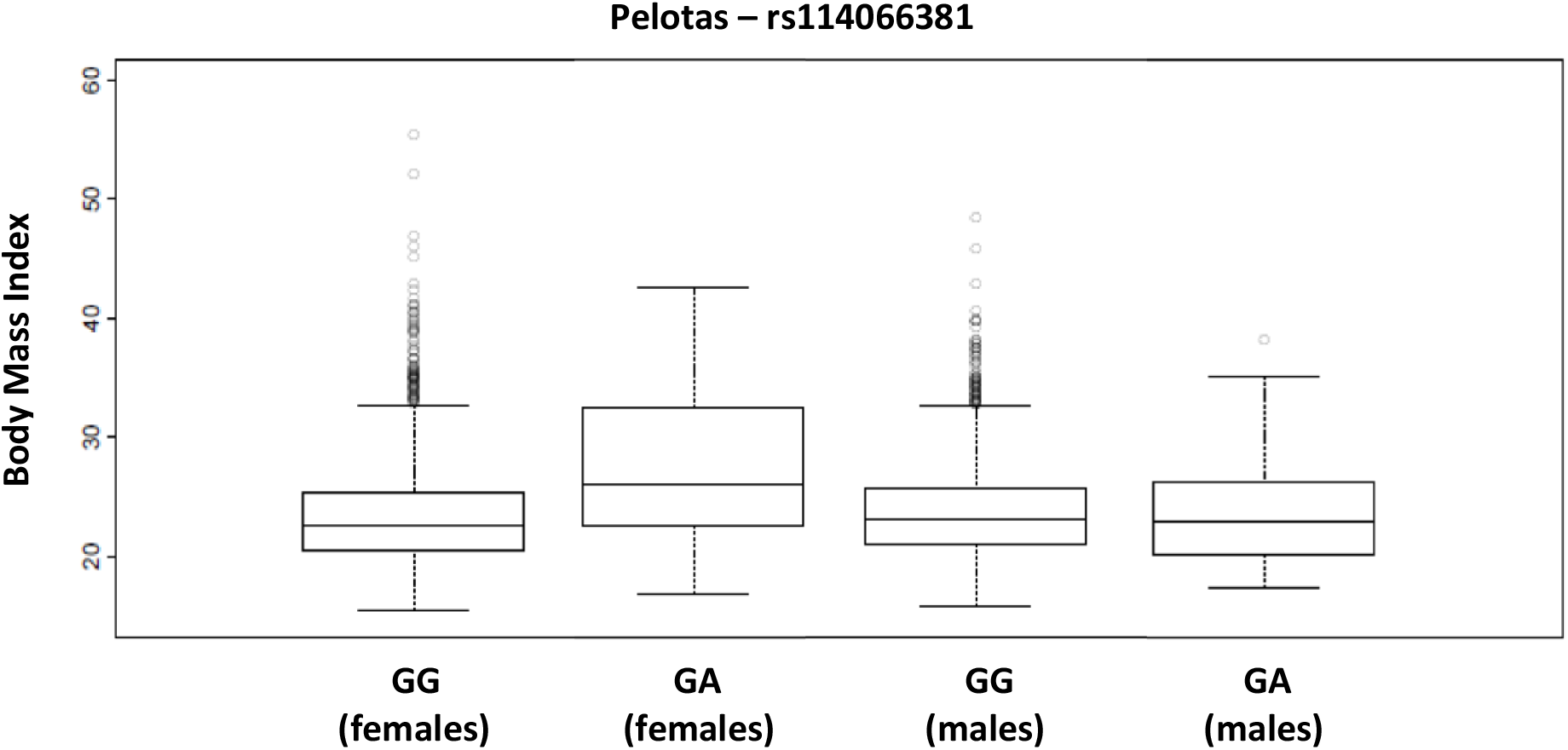
Body Mass Index (BMI) in females and males’ adults from Pelotas cohort, according to their genotypes in the SNP rs114066381. The increase of BMI associated with the rs114066381-A is observed in females (p-value =0.0008), but not in males (p-value =0.5397)

**Table 2.**
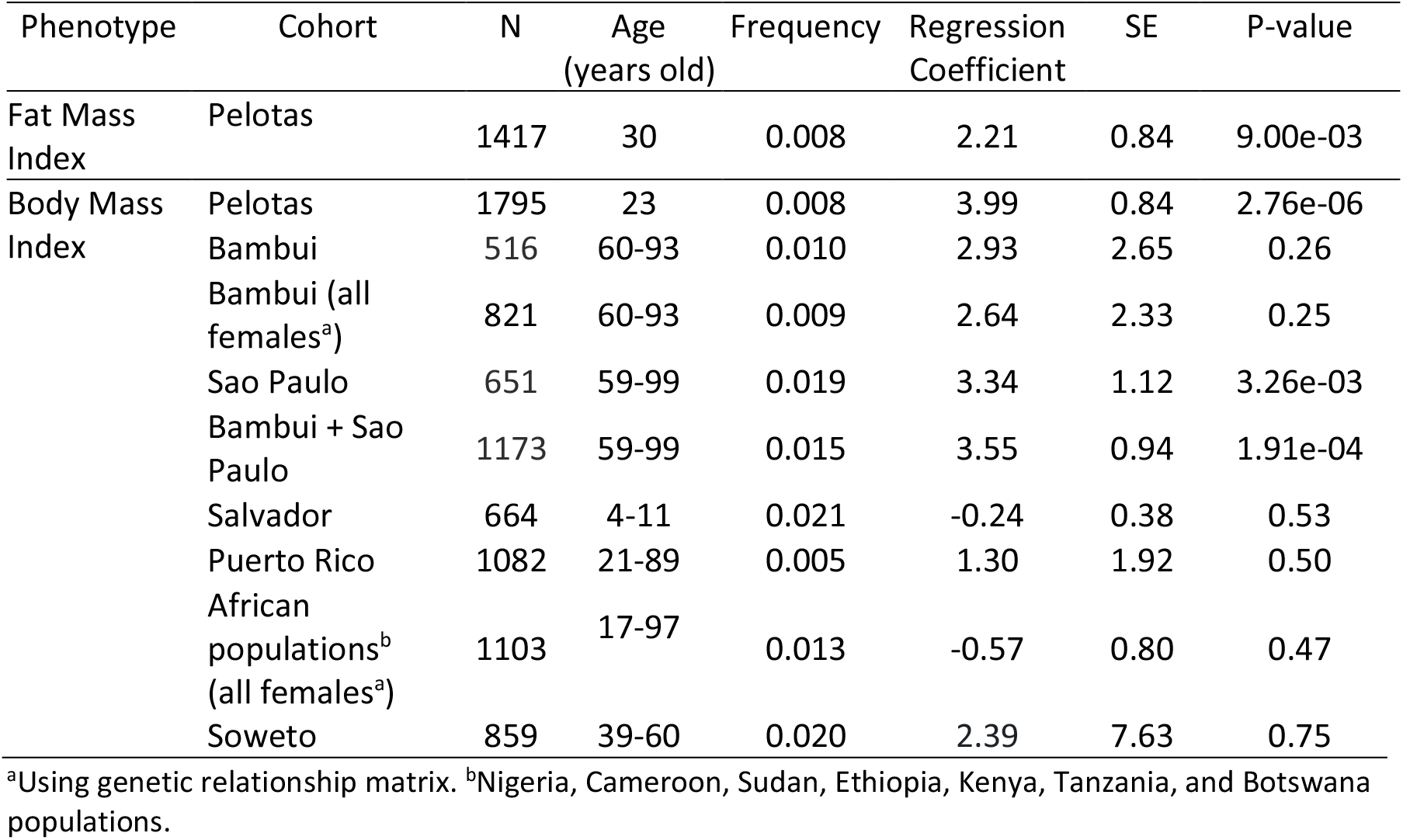
Association of rs114066381-A with Body Mass Index and Fat Mass Index in unrelated females of the Pelotas cohort and replications.

| Phenotype | Cohort | N | Age<br>(years old) | Frequency | Regression<br>Coefficient | SE | P-value |
| --- | --- | --- | --- | --- | --- | --- | --- |
| Fat Mass<br>Index | Pelotas | 1417 | 30 | 0.008 | 2.21 | 0.84 | 9.00e-03 |
| Body Mass<br>Index | Pelotas | 1795 | 23 | 0.008 | 3.99 | 0.84 | 2.76e-06 |
|  | Bambui | 516 | 60-93 | 0.010 | 2.93 | 2.65 | 0.26 |
|  | Bambui (all<br>females<sup>a</sup>) | 821 | 60-93 | 0.009 | 2.64 | 2.33 | 0.25 |
|  | Sao Paulo | 651 | 59-99 | 0.019 | 3.34 | 1.12 | 3.26e-03 |
|  | Bambui + Sao<br>Paulo | 1173 | 59-99 | 0.015 | 3.55 | 0.94 | 1.91e-04 |
|  | Salvador | 664 | 4-11 | 0.021 | -0.24 | 0.38 | 0.53 |
|  | Puerto Rico | 1082 | 21-89 | 0.005 | 1.30 | 1.92 | 0.50 |
|  | African<br>populations<sup>b</sup><br>(all females<sup>a</sup>) | 1103 | 17-97 | 0.013 | -0.57 | 0.80 | 0.47 |
|  | Soweto | 859 | 39-60 | 0.020 | 2.39 | 7.63 | 0.75 |

<sup>a</sup>Using genetic relationship matrix. <sup>b</sup>Nigeria, Cameroon, Sudan, Ethiopia, Kenya, Tanzania, and Botswana populations.

rs114066381 is 2 kb from a CTCF-binding site (Zerbino et al. 2018), but no evidence of transcription regulation is shown by RNA-seq (ENCODE Project Consortium 2011). Besides, this genomic region contains binding sites for the histone-interacting proteins KAP1 and SETDB1, as reported by ChIP-seq data (HaploReg v4.1, Ward and Kellis 2016; ENCODE Project Consortium 2011; Figure S7). However, there is no evidence in the literature that the region acts as an enhancer in vivo. This genomic region is primate-specific, being absent from the genome of other vertebrates (UCSC Genome Browser 2013, Kent et al. 2002, Figure S8, S9). The derived allele A is very rare in Europeans, but has frequencies of ~3% in West Africans (Table S6).

We confirmed the rs114066381 female-specific association using the fat mass index (a more direct measurement of adiposity), measured by Dual-energy X-ray absorptiometry (DXA), seven years after the measurement of BMI on the same individuals (beta= 2.21 kg of fat/m^2^ per allele, 95%CI: 0.55-3.88, p=9×10-3, Table 2). We replicated with 89% of power the association in older adult females from São Paulo (Brazil) (SABE cohort, Barbosa et al. 2005) and in the merged dataset from São Paulo and Bambuí (power=99%, Table 2). We tested but did not observe significant association (Table 2) in females from Bambuí cohort alone (power=53%), in girls from Salvador (power=9%), in 1082 women from a breast cancer Puerto Rico cohort (power=13%), in 859 women from Soweto (South Africa, power=69%), and in 1085 women from different ethnic groups settled in rural Africa (power=9%). However, a meta-analysis synthesizing the seven effect sizes showed a positive association between rs114066381-A allele and BMI in females (Figure 4) both considering all effects together or only the effects obtained with admixed populations. Remarkably, while in Pelotas and São Paulo the frequency of rs114066381-A allele is 1.1%, its frequency increases in overweight (1.2%) and obese (1.98%) women, and attains 9% among morbidly obese women. The same pattern is also observed in Bambuí and Soweto populations but not in Puerto Rico, in which the frequency of rs114066381-A allele is higher only for obese women (Table S10). We observed that the BMI distribution of rural Africa and Salvador are very different compared to the other populations: while in Pelotas, Bambuí, São Paulo, Puerto Rico, and Soweto, 27% of individuals have BMI greater than 25 kg/m^2^, in Salvador and rural Africa, 11% of individuals fall in this category (and only 0.2% of individuals are morbidly obese in both cohorts, contrasting to more than 1% in the other populations). Moreover, 23% of the individuals of rural Africa are underweight (BMI < 18.5 kg/m^2^).

**Figure 4.**
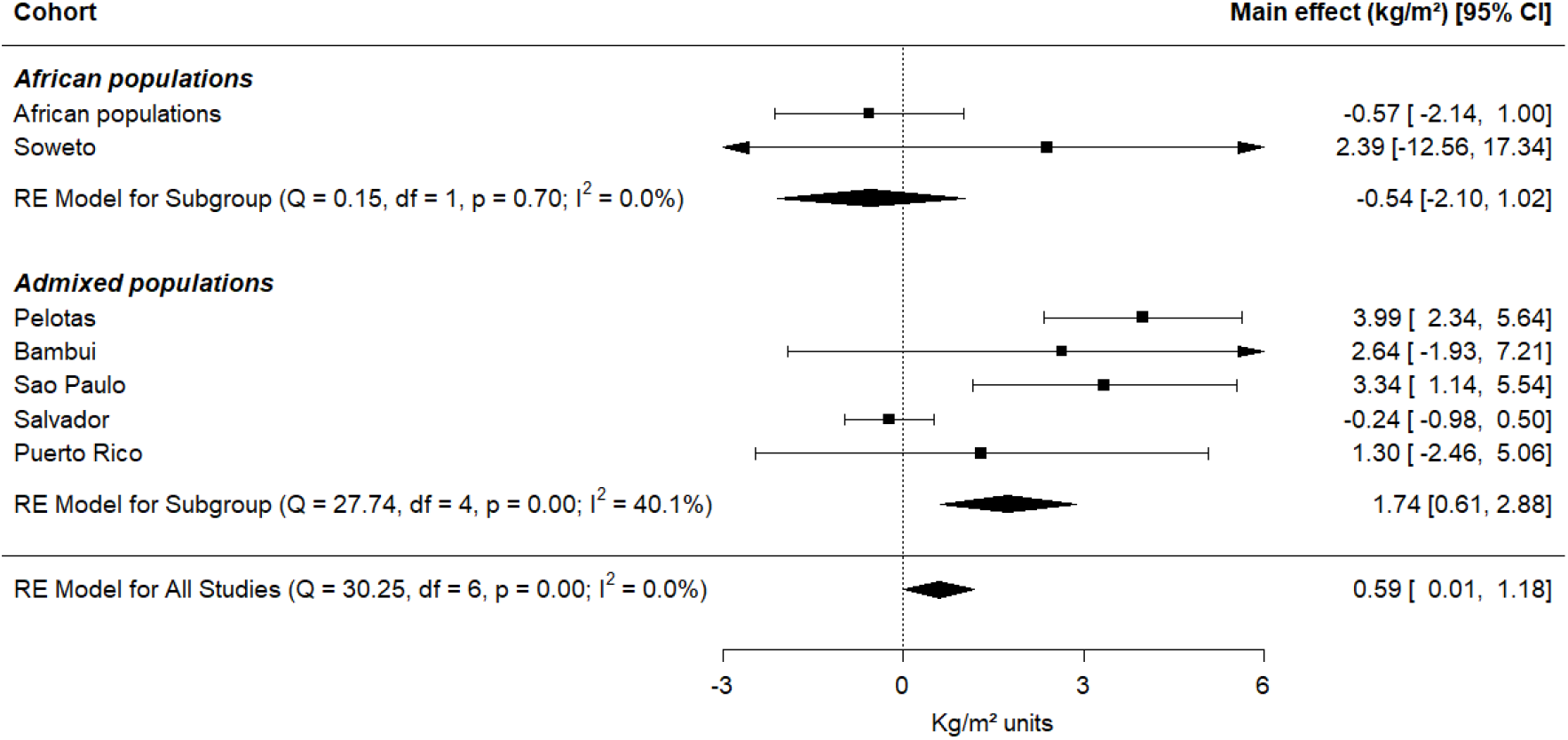
Forest plots from the meta-analysis synthesizing association results between rs114066381 and BMI from seven populations. Effect size [95% confidence interval (CI)] in each individual study, subgroups of African populations and admixed populations, and combining all populations.

### Power estimation and Replication of previous GWAS hits

We estimated the statistical power to detect associations for 216 BMI GWAS-catalog-hits on the three EPIGEN cohorts, conditioning on the effect sizes reported in kg/m^2^ units on the GWAS-Catalog (took as a population parameter), the BMI distribution, as well as the number of studied individuals and the allele frequencies in each of the EPIGEN cohorts (Supplementary Data). These 216 GWAS-Catalog-hits were selected because, in the context of the high heterogeneity of data stored in the GWAS-Catalog, their effect sizes (linear regression coefficient) were unambiguously associated with a specific allele and were consistently reported in units of kg/m^2^. Out of the 216 hits, one hundred and eighty-nine were observed in adults, 4 in children, and 23 in both, in individuals with predominant European-ancestry.

Based on the mean statistical power of the 216 GWAS-Catalog-hits (Supplementary Data), and assuming that these SNPs (and their regression coefficient) are part of the genetic architecture of BMI in the Brazilian cohorts, we would expect to observe 24 SNPs out of 216 (average power=11%) associated in Salvador children, 22 SNPs (average power=10%) in Pelotas young adults and 15 SNPs (average power=7%) in Bambuí older adults (Figure S10); however we observed 0, 20, and 8 significantly associated SNPs on the replication analysis for the three cohorts, respectively (Table S11). Specifically, in Pelotas, we confirmed the association for the six *FTO* SNPs included in our analysis (rs9930333, rs62033400, rs8050136, rs3751812, rs1558902, and rs9939609).

## DISCUSSION

Leveraging admixture in Brazil, we used genome-wide data of three population-based cohorts to find loci associated with BMI through admixture mapping followed by fine-mapping. While a limitation of our study is the relatively small number of individuals respect to GWAS standards, it has important characteristics absent in most studies. First, it relies on population-based cohorts that better capture the phenotypic variation of populations, but are rarely considered in genetic studies (Lasky-Su et al. 2008; Manolio et al. 2006). Second, it presents the first admixture mapping performed in a South American admixed population, releasing the respective masterscripts to perform this kind of studies in other populations (http://www.ldgh.com.br/scientificworkflow/master_scripts.php). Third, it is one of the few studies that explore the genetic architecture of BMI in three different age stages: children, young adults and older adults. Finally, in the context of under-representation of non-European populations in genome-wide association studies, we analyzed populations of South America with African and Native American ancestry.

Because none of the six new candidate SNPs to influence BMI reported in this study (Tables S7 and S9) are in LD with previous 389 GWAS-Catalog-hits of BMI (r^2^<0.022), we conclude that, by studying the admixed populations of Salvador, Pelotas and Bambuí, we have contributed to expand the catalog of SNPs of the global genetic architecture of BMI.

### Factors supporting the female-specific effect of rs114066381 on adiposity

First, we replicated the female-specific association in an independent admixed cohort from São Paulo and confirmed its association by a meta-analysis. Second, while our initial female-specific association with BMI and fat-mass index is based on imputed genotypes, replication in the São Paulo cohort is based on whole genome data. Third, in the discovery cohort, we not only observed a strong association with BMI, but also with fat mass index measured with DXA, which is a more direct measure of adiposity. Importantly, the fat mass index was collected seven years later than BMI in the same individuals. Fourth, rs114066381 is located in a potential regulatory region, which makes a biological role for rs114066381 plausible. With respect to the pattern of linkage disequilibrium, rs114066381 presents a r^2^=0.5 with three SNPs in linkage disequilibrium with each other (r^2^=1) mapped in near regulatory regions (in sensu HaploREG v4.1 in Ward and Kellis 2016, Figure S11). In Bambuí the r^2^ of rs114066381 with the same three SNPs varies between 0.3 and 0.7 (Figure S11). These results suggest a specific role for this SNP. rs114066381 is ~300 kb from rs7335631, previously associated with “Fat distribution in HIV” (Irvin et al. 2011), but there is no (r^2^<0.001) LD between these two SNPs in any of our three Brazilian populations.

The discovery size effect for rs114066381 (beta=3.99±0.847 kg/m^2^) is one of the highest observed for BMI, considering both sex. The size effects suggested by both meta-analyses, Figure 4) are also high considering the distribution of BMI-effect sizes. According with GWAS-Catalog (October 2019), the range of estimated beta in kg/m^2^ for BMI-hits is 0.013 - 4.119 with an average of 0.054 kg/m^2^.

The female-specific association for rs114066381 is observed in the following context: out of 833 hits reported in GWAS-Catalog as associated with BMI with beta reported in kg/m^2^ (October 2019) independently of sex, 229 are female-specific associations (beta range: 0.009 - 0.484, beta mean: 0.025) and 134 male-specific [beta range: 0.013 - 0.095, beta mean: 0.025]. Even if the mean effect sizes are similar in men and women, the effect size distribution of women shows a tail of higher beta values, which suggest that there are more genetic variants predisposing to higher BMI in women than in men. Our finding is paradigmatic of this context: in adult females of Southern Brazil, rs114066381 alone explains a similar portion of the variance of BMI (r^2^ range for Pelotas, Bambuí, and São Paulo cohorts: 0.008 to 0.044) as the entire set of 97 GWAS-hits recently reported by Locke et al. (2015).

### Methodological issues

Inferences of continental local chromosome ancestry based on genome-wide data are more uncertain than inferences about individual or population ancestry. For this reason, we report consensus admixture mapping hits, which means that they correspond to genomic regions whose ancestry is associated with BMI for local ancestry inferred both with PCAdmix (Brisbin et al. 2012) and RFMix (Maples et al. 2013). Notably, the five consensus admixture mapping hits not only match for the chromosome band, but more specifically, correspond to overlapping genomic regions. In general, to avoid spurious results in admixture mapping studies, we suggest to validate the AM results using at least two different methods for local ancestry inference.

### Replication of other GWAS hits

We replicated 28 of the 216 associations reported for SNPs in previous GWAS, mostly performed in adults of European ancestry, with the Pelotas cohort presenting not only the largest rate of replication (20/216) (Table S11), but also a very good concordance between the observed (20) and expected (22) number of replications. This is consistent with: (1) the larger size of Pelotas cohort; (2) the relative lower socioeconomic status of the Salvador cohort adds a layer of complexity to the definition of the genetic architecture of BMI, respect to GWAS in predominantly European populations with different socio-economic background. (3) the age-dependence of the genetic architecture of BMI, and the fact that most GWAS of BMI were performed in adults and in Pelotas BMI was also measured in young adults, while in the Salvador and Bambuí cohorts, BMI was measured in children and older adults, respectively. Indeed, Lasky-Su et al. (2008) showed how age-dependent effects can be an important and misjudged cause of non-replication.

In conclusion, we performed three admixture mapping/fine-mapping for BMI, and tested the association of GWAS-Catalog-hits in three Brazilian population-based cohorts. These cohorts present different levels of African ancestry, socioeconomic background, and different ages (children, young adults and older adults), and therefore, our study contributes to better define the global and age-dependent genetic architecture of BMI. We provide a list of six candidate SNPs associated with African or European ancestry that are associated with BMI. More importantly, our admixture/fine mapping in Brazilians reveals a West African associated potential regulatory variant (rs114066381), with a female-specific effect on BMI, which seems to be particularly important for the development of morbid obesity. Altogether, our results show that the study of South American admixed populations, as well as other populations worldwide (Chen et al. 2017; Granot-Hershkovitz et al. 2018; Salinas et al. 2016) are a source of novel non-European associated variants with considerable effect size and that may explain in non-European populations an important portion of the current *missing heritability*.

## MATERIALS AND METHODS

### Study populations and genotyping

The Salvador-SCAALA is a cohort study comprising 1,445 children aged 4-11 years in 2005, living in Salvador (Figure 1), a city of 2.8 million inhabitants in northeast Brazil (Barreto et al. 2006). This population is part of an earlier observational study and represent the population without sanitation in Salvador. BMI was measured in 2005, when children were 4 to 7 years old.

The 1982 Pelotas birth cohort study was conducted in Pelotas, a city in southern Brazil, with 340,000 inhabitants (Figure 1). Throughout 1982, the three maternity hospitals in the city were visited daily and births were recorded, corresponding to 99.2% of all births in the city. Of these, the 5,914 liveborn infants whose families lived in the urban area constituted the original cohort (Victora & Barros 2006). BMI was measured at 2004 and 2005, when individuals were 23 years old. The 2012-2013 follow-up of the cohort included a detailed assessment of body composition. Trained technicians measured participants’ body fat, lean and bone-mineral masses at the Pelotas cohort research clinic using dual-energy X-ray absorptiometry (DXA; GE Lunar Prodigy densitometer) in a full-body scan. We calculated fat mass index by dividing the adjusted fat mass (kg) by height (m) squared.

The Bambuí cohort study of ageing is ongoing in Bambuí, a city of approximately 15,000 inhabitants, in Minas Gerais State in southeast of Brazil (Figure 1). The population eligible for this cohort consisted of all residents aged 60 years and over on January 1997, who were identified from a complete census in the city. From 1,742 eligible residents, 1,606 constituted the original cohort (Lima-Costa et al. 2011; Lima-Costa et al. 2015a, 2015b). BMI was measured in 1997, when individuals were between 60 to 93 years old.

The EPIGEN-Brazil initiative genotyped 1,222 individuals studied for BMI from the Salvador cohort, 3,628 from the Pelotas cohort, and 1,342 from the Bambuí cohort, using the Illumina (San Diego, CA, US) Omni 2.5M array as detailed in Kehdy et al. (2015).

For the three cohorts, BMI was calculated as weight (kg) divided by squared height (meters) and these measurements were taken by trained research staff. Potential confounding variables like sex, age, and different socioeconomic characteristics were collected for the three cohorts (Table S1). We categorized socioeconomic variables in each cohort as follows. For Salvador, two classes were defined on the basis of maternal schooling: (1) complete elementary school or less (8 years or less) and (2) at least incomplete high school (> 8 years). For Pelotas, two classes were defined based on family income, expressed as minimum wages (equivalent to USD 45,00 in 1982): (1) as lower than one minimum wage, (2) equal or higher to one minimum wage. For Bambuí, two classes were defined based on schooling: (1) incomplete primary school or less (<4 years), and (2) at least complete primary school (> 4 year); additionally, monthly household income per capita was categorized into: (1) lower and (2) equal or higher to the median value (equivalent to USD 180.00 in 1997).

The EPIGEN protocol was approved by Brazil’s national research ethics committee (CONEP, resolution number 15895, Brasília). Kehdy et al (2015) estimated the proportions of African, European, and Native American ancestries for each individual of each cohort (Figure 1, Table S1) using the software ADMIXTURE (Alexander et al. 2009) and we used those estimates in the present study.

### Kinship coefficients

Kinship coefficients between pairs of individuals were estimated using the REAP method (Thornton et al. 2012), which is appropriate for admixed populations. For Salvador and Pelotas we eliminated 63 and 83 individuals, respectively, that had relatives in the cohort (kinship coefficient higher than 0.1) using a network-based approach that aims to eliminate the smallest possible number of individuals (Kehdy et al. 2015). The Bambuí cohort, has 516 (36%) individuals with relatives in the cohort. Thus, for Bambuí we identified families with a categorical variable, and used robust variance estimators to correct results by family structure. The final dataset used for the analysis included 1,222 individuals from Salvador, 3,628 individuals from Pelotas and 1,342 individuals from Bambuí.

### Phasing and local ancestry inference by PCAdmix and RFMix software

We phased our datasets using the software SHAPEIT2 (Delaneau et al. 2011), as detailed in (Kehdy et al. 2015). We used two methods for local ancestry inference, implemented in the software PCAdmix (Brisbin et al. 2012) and RFMix (Maples et al. 2013).

PCAdmix inferences were performed as in Kehdy et al. (2015). For RFmix inferences, we used as parental populations: Europeans (CEU and IBS) from 1000 Genomes Project (1000G, 2015); Africans from Botswana (Crawford et al. 2017), Ghana (from National Cancer Institute (NCI) Survey of Prostate Cancer in Accra, Gouveia et al. 2019) and GWD from 1000G (2015); and eight Native American populations (Quechuas, Ashaninkas, Matsiguengas, Aymaras from Tarazona-Santos group and Matsiguengas, Qeros, Uros and Moches from Harris et al. 2018 and from Tarazona-Santos group). For X chromosome inferences, we used as parental populations the same Europeans and Native Americans populations (the four populations from Tarazona-Santos group) referred above, and Africans (LWK and YRI) from 1000G. RFMix uses a conditional random field parameterized by random forests trained on reference panels, learning from the admixed samples to autocorrect phasing errors and improve local ancestry inferences (Maples et al. 2013). To run RFMix we fixed the number of generations since the admixture event (parameter -*G*) in 20 (~500 years) and the number of trees to generate per random forest (parameter -*t*) in 500. Inferences were performed in window lengths (parameter -*w*) of 0.2 cM. All other parameters present in RFMix were set as default. For PCAdmix and RFMix results, we considered only the windows which ancestry was inferred with a posterior probability > 0.90.

### Relationship between BMI and individual ancestry

To assess the effect of individual admixture (European, African and Native American) on BMI, we used a multiple linear regression including age (in Salvador and Bambuí cohorts only), sex, and socioeconomic variables as covariates. We performed a generalized linear model with Gaussian distribution for all three cohorts. In the case of Bambuí we additionally used the Sandwich R package (Robust Covariance Matrix Estimators, Zeileis 2004) to correct the estimations by family clusters (assigning to each individual a categorical variable that represents his/her family). The variables included in the final regression model were selected by performing a forward stepwise method using the Akaike information criterion (AIC). The final model included all the covariables considered above. All statistical analyses were performed using R platform (R Core Team, 2014).

### Admixture mapping

We performed admixture mapping using inferences from two methods: PCAdmix and RFMix. Only consensus hits for both methods are reported in this study. We tested the association between BMI and each local ancestry (African, European, and Native American) across the genome using linear regression models. The regressions were adjusted by age (Salvador and Bambuí), sex, socioeconomic variables, and genome-wide African ancestry. For Bambuí, we included the correction for family structure. We used an additive model that considers the number of inferred African, European or Native American ancestry copies (0, 1 or 2) carried by an individual for each window. Because we found an association between individual African ancestry and BMI in females in Pelotas, we performed a stratified analysis for each sex in this cohort. We used PLINK software (Purcell et al. 2007) for regression analysis for Salvador and Pelotas cohorts. For Bambuí, we used a model-robust standard error estimators implemented in the Sandwich package in R environment (Zeileis 2004).

To establish a significance threshold for the admixture mapping, accounting both for multiple testing and linkage disequilibrium due to admixture, we estimated the effective number of tests (ENT) for each chromosome for each individual as in Shriner et al. (2011). The method fits an autoregressive model to the vector of local ancestries (0, 1, or 2 chromosomes of given ancestry) and evaluates the spectral density at frequency zero with the package code for R. We used the estimated ENT to obtain a Bonferroni p-value threshold for significance as 0.05 divided by ENT. The number of tests was estimated using PCAdmix inferences. Genome-wide p-values thresholds were obtained for each ancestry in each dataset (Table S5) and were within the range 10e-04 - 10e-06. We conservatively used the same genome-wide thresholds to identify admixture mapping hits for X-chromosome. In our approach, we compared the significant admixture mapping peaks of each chromosome obtained using PCAdmix with the significant admixture mapping peaks obtained using RFMix inferences for the same chromosomes (using p-values thresholds from Table S2 multiplied by 10), and, considered a consensus significant peak the ones mapped to the same chromosomal region (not only the same chromosome bands) using both inferences. Only these consensus significant admixture mapping peaks were followed-up for fine-mapping. For chromosome windows showing the most significant admixture mapping hits, we used analysis of variance to test if the BMI means differ between individuals carrying 0, 1 or 2 copies of chromosome windows of a specific ancestry.

### Imputation, Fine-mapping, Annotation

Fine-mapping of significant admixture mapping peaks was performed using both genotyped and imputed SNPs. We used an EPIGEN-Brazil dataset imputed with IMPUTE2 (Howie et al. 2009), focusing on ±1 Mb centered in the most significant window of each admixture mapping hit (based on PCAdmix). We imputed our dataset with a reference panel that merged the public reference panel data from 1000G and 270 individuals from EPIGEN (90 of each cohort) genotyped for 4.3 million SNPs, and considered only SNPs imputed with an info score quality metric > 0.8 (Magalhães et al. 2018).

Genotyped and imputed SNP were tested for association with BMI using the same linear regressions models used for admixture mapping. We excluded SNPs with minor allele frequency < 0.005 for these analyses. We considered significant, the associations with p-values less than or equal to the ones obtained for the admixture mapping peaks and suggestively significant those SNPs with a p-value higher than the ones obtained for the admixture mapping peaks but not more than one unit of -log (p-value). Fine-mapping results were plotted using the LocusZoom tool (Pruim et al. 2010). We used ANNOVAR (Wang et al. 2010) to annotate associated genetic variants. We estimated the linkage disequilibrium statistics (r^2^, Hill 1968) on phased data using the software Haploview (Barrett et al. 2005).

A flowchart summarizing the study design is shown in Figure S12. Both the flowchart as well as the bioinformatics scripts used for the analyses are available in the EPIGEN-Brazil Project Scientific Workflow (http://www.ldgh.com.br/scientificworkflow, Magalhães et al. 2018).

### Replication cohorts and meta-analysis

We tested for replication of the association of rs114066381-A with BMI in other four cohorts, as follows: (i) in whole-genome data from 651 unrelated females from São Paulo, Brazil, the SABE (Health, Well being and Aging) study (Barbosa et al. 2005; Naslavsky et al. 2017). Whole-genome sequencing data were generated at Human Longevity Inc. (HLI, San Diego, California) with a mean target coverage of 30x. More information about the sequencing and bioinformatics pipelines used can be found in Telenti et al. (2016). Linear regression was adjusted for age, education level, SES, and African Ancestry proportion; (ii) imputed data from 1,082 women from Puerto Rico (547 non-cancer controls and 535 cases). Linear regression was adjusted for age, education level (as a measure of SES), African Ancestry proportion, and breast cancer diagnosis; (iii) genotyped and imputed data from 1,103 adult women (age >= 18 years) from Nigeria, Cameroon, Sudan, Ethiopia, Kenya, Tanzania, and Botswana (Crawford et al. 2017). All individuals were genotyped on either the Illumina 1M-Duo BeadChip array or the Illumina 5M-Omni array. The polymorphism is directly typed on the Illumina 5M Omni array and imputed for individuals typed on the Illumina 1M-Duo BeadChip array. Association tests were performed using a linear-mixed model, as implemented by the Genome-wide Complex Trait Analysis (GCTA) software. In this model, age was modeled as a fixed effect and the kinship matrix was used for the random effects term; (iv) imputed data from 859 women from Soweto, South Africa (Ali et al. 2018). Linear regression was adjusted for age and SES.

We used the package metafor from R code (Viechtbauer 2010) to perform the meta-analysis to synthesize the effects from seven different populations. We used Random-Effects model with Hedges method.

### DANCE Analysis

We obtained a graph representation of the genetic architecture of BMI using the DANCE web tool (Disease-ANCEstry Networks, http://gilderlanio.pythonanywhere.com/home, Araújo et al. 2016), a network-based computational approach to integrate, summarize and visualize GWAS-Catalog-hits (https://www.ebi.ac.uk/gwas/) and 1000G allele frequency information for different populations. We also used DANCE to visualize the genetic architecture of associated genes.

### Statistical power estimation

Power estimation was performed separately for each EPIGEN cohort, according to the specific BMI distribution (Table S1). SNPs associated with BMI in the GWAS-Catalog were extracted using the keyword “body mass index”, and those with p-values > 9e-05 were filtered out. From the 2205 SNPs reported on 61 published studies, we kept 216 SNPs with effect size unambiguously associated to a specific allele and reported as a regression coefficient expressed in kg/m^2^ from cross-sectional studies, genotyped or imputed in our database. We calculated the statistical power using the latter effect size (regression coefficient) values, and allele frequency and number of individuals from each EPIGEN cohort (Table S1). The type I error rate was set at α = 0.00023. All power estimates were calculated with QUANTO v1.2.4 program (Gauderman & Morrison 2006), assuming an additive genetic model with independent individuals.

### Replication analysis of previous GWAS hits

To test the association of previous BMI GWAS hits we used the regression model used in fine-mapping for all the selected 216 SNPs in the three Brazilian cohorts. P-values were adjusted considering 216 independent tests using the Benjamini-Hochberg correction (Benjamini & Hochberg 1995).

### Genomic in-silico analyses

The search for candidate regulatory SNPs was performed in May 25, 2018 using HaploReg v4.1 database (http://archive.broadinstitute.org/mammals/haploreg/haploreg.php, Ward and Kellis 2016), Ensembl (https://grch37.ensembl.org/, Zerbino et. 2018), and RegulomeDB (http://www.regulomedb.org/, Boyle et al. 2012). ChIP-seq data was provided by ENCODE Project Consortium (2011), available at HaploReg v4.1 database. We used MULTIZ from UCSC Genome Browser, Blanchette et al. 2004) to determine whether there are homologous of the region where SNP rs114066381 in the genomes of 30 other placental mammals. For that purpose, we used as query a human genomic sequence with 2001 nucleotides centered in polymorphism rs114066381 (human genome version hg38 chr13: 30057599-30059599). We used TimeTree to generate the phylogenetic tree (Kumar et al. 2017) providing species names as query (Figure S9 contains the correspondence between common names as provided by MULTIZ and the species names used to generate the tree).

## ACKNOWLEDGEMENTS

For analyses, we used the Sagarana cluster (from Centro de Laboratórios Multiusuários do Instituto de Ciências Biológicas-Universidade Federal de Minas Gerais). We thank Miguel Ortega for help in the use of Sagarana, Ms. Evelyn Tay at University of Ghana Medical School (Accra, Ghana) for managing the study, and Ms. Marcelle Bartholomeu and Ms. Àlex Teixeira for technical support. The EPIGEN-Brazil Initiative is funded by the Brazilian Ministry of Health (Department of Science and Technology from the Secretaria de Ciência, Tecnologia e Insumos Estratégicos) through Financiadora de Estudos e Projetos. The EPIGEN-Brazil investigators received funding from the Brazilian Ministry of Education (CAPES Agency), Brazilian National Research Council (CNPq), the Minas Gerais State Agency for Support of Research (FAPEMIG) and TWAS-CNPq PhD fellow.

