## Supplementary Tables and Figures for "Admixture/fine-mapping in Brazilians reveals a West African associated potential regulatory variant (rs114066381) with a strong female-specific effect on body mass- and fat mass-indexes" (from another version of this preprint: version 1, 2019-11-14)

**Table S1. Demographic characteristics and Body Mass Index (BMI) of the studied population-based cohorts (Salvador, Pelotas and Bambui).**

| Characteristics | Cohorts | | |
|---|---|---|---|
| | Salvador children (n=1222) | Pelotas young adults (n=3628) | Bambui older adults (n=1342) |
| **Sex** | | | |
| Female | 664 (54.3%) | 1798 (49.6 %) | 821 (61.2%) |
| Male | 558 (45.7%) | 1830 (50.4%) | 521 (38.8%) |
| **Age (years)**[a] | 6.83 ± 1.67 (4-11) | 23 | 68.8 ±7.0 (60-93) |
| **BMI (kg/m²)**[a] | 15.54 (±2.05) | 23.72 (±4.38) | 25.12 (±4.92) |
| **BMI classification** [b] | | | |
| Underweight | 0 \| 37(3.0%) | 214 (5.9%) | 94 (7.0%) |
| Normal | 1091(89.3%) \| 1049(85.9%) | 2348 (64.7%) | 597 (44.5%) |
| Overweight | 78 (6.4%) \| 108 (8.8%) | 764 (21.1%) | 479 (35.7%) |
| Obese | 53 (4.3%) \| 26 (2.1%) | 278 (7.6%) | 159 (11.8%) |
| Morbid Obese | 0 \| 2 (0.2%) | 24 (0.7%) | 13 (1.0%) |
| **Educational level**[c] | | | |
| <= Elementary school | 668 (54.7%) | - | - |
| >= High school | 554 (45.3%) | - | - |
| <4 years | - | - | 860 (64.4%) |
| >= 4 years | - | - | 389 (27.9%) |
| **Monthly income**[d] | | | |
| <= 1MW | - | 738 (20.3%) | - |
| > 1MW | - | 2890 (79.7%) | - |
| < USD 180 | - | - | 629 (46.9%) |
| >= USD 180 | - | - | 713 (53.1%) |
| **Genomic ancestry**[a] | | | |
| African | 0.51 (±0.14) | 0.16 (±0.19) | 0.14 (±0.15) |
| European | 0.43 (±0.13) | 0.76 (±0.20) | 0.79 (±0.16) |
| Native American | 0.06 (±0.03) | 0.08 (±0.05) | 0.07 (±0.04) |
| **X-Chromosome ancestry**[a] | | | |
| African | 0.58 (±0.24) | 0.19 (±0.25) | 0.16 (±0.19) |
| European | 0.30 (±0.22) | 0.67 (±0.27) | 0.69 (±0.22) |
| Native American | 0.12(±0.10) | 0.14 (±0.12) | 0.15 (±0.12) |

[a]Mean (±SD); [b]BMI was classified according to the WHO guidelines: for children we used the Z-scores adjusted by sex and age (<5yo, >=5yo). For adults: Underweight (BMI<18.5), Normal weight (>=18.5 and <25), Overweight (>=25 and <30), Obese (>=30 and <=40) , and Morbide obese (>40). Because WHO does not report a cutoff for morbid obesity in children, we also reported in percentiles adjusted by sex and age (<5yo, >=5yo) following CDC quidelines.; [c]Mother educational level for Salvador; [d]MW = Minimum wage (USD 45 in 1982) in Pelotas cohort, the value of USD 180 in Bambui corresponds to the median of income in Bambui cohort.

**Table S2. Multiple linear regression analyses of BMI on African, European, and Native American ancestries for children from Salvador cohort (n=1222).**

| Variables | African | | European | | Native American | |
|---|---|---|---|---|---|---|
| | β | p-value | β | p-value | β | p-value |
| Age (years) | 0.22 | **4e-10** | 0.22 | **4e-10** | 0.22 | **4e-10** |
| Sex Male | -0.22 | 0.052 | -0.22 | 0.053 | -0.22 | 0.055 |
| Mother educational level | | | | | | |
| >= High school | 0.47 | **5e-05** | 0.47 | **5e-05** | 0.48 | **3e-05** |
| Genomic ancestry[a] | -0.03 | 0.422 | 0.04 | 0.411 | 0.04 | 0.850 |
| Interaction: ancestry*male | 0.06 | 0.473 | -0.03 | 0.755 | -0.63 | 0.093 |
| Genomic ancestry[b] | | | | | | |
| 0.21-0.40[c] \|> 0.20[d] | 0.53 | 0.274 | -0.50 | 0.137 | 0.96 | 0.407 |
| 0.41-0.60[c] | 0.10 | 0.834 | -0.55 | 0.096 | - | - |
| 0.61-0.80[c] | 0.24 | 0.618 | -0.09 | 0.805 | - | - |
| > 0.80[c] | 1.05 | 0.169 | -1.45 | 0.317 | - | - |

Multiple linear regression adjusted by age, sex, mother educational level, and ancestry ($\beta_0$ for African ancestry = 14.11, $\beta_0$ for European ancestry = 13.79, $\beta_0$ for Native American ancestry = 13.90 kg/m2); [a]Ancestry value in tenths; [b]Ancestry in classes of 0.20 units of ancestry, where reference on regression analysis is the class of ancestry 0-0.20; [c]For African and European ancestries there are five intervals, [d]For Native American ancestry there are two intervals. Values in bold indicate p-value < 0.05.

**Table S3. Multiple linear regression analyses of BMI on African, European, and Native American ancestries for young adults from Pelotas cohort (n=3628).**

| Dataset | Variables | | African | | European | | Native American | |
|---|---|---|---|---|---|---|---|---|
| | | | β | p-value | β | p-value | β | p-value |
| ALL | Sex | Male | 0.73 | **1e-04** | -2.50 | **1e-05** | 1.09 | **2e-04** |
| | Income | > 1MW | 0.36 | 0.052 | 0.38 | **0.042** | 0.27 | 0.138 |
| | Genomic ancestry[a] | | 0.29 | **9e-08** | -0.31 | **2e-09** | 0.85 | **1e-04** |
| | Interaction: ancestry*male | | -0.33 | **2e-05** | 0.36 | **9e-07** | -1.10 | **4e-04** |
| | Genomic ancestry[b] | | | | | | | |
| | | 0.21-0.40[c] \|>0.20[d] | 0.59 | 0.105 | 1.04 | 0.225 | -0.21 | 0.769 |
| | | 0.41-0.60[c] | 2.06 | **5.E-08** | 0.56 | 0.505 | - | - |
| | | 0.61-0.80[c] | 1.24 | **0.006** | -0.54 | 0.501 | - | - |
| | | > 0.80[c] | -0.70 | 0.599 | -0.89 | 0.256 | - | - |
| FEMALE | Income | > 1MW | -0.06 | 0.819 | -0.01 | 0.974 | -0.23 | 0.390 |
| | Genomic ancestry[a] | | 0.27 | **3e-06** | -0.29 | **1e-07** | 0.80 | **8e-04** |
| | Genomic ancestry[b] | | | | | | | |
| | | 0.21-0.40[c] \|>0.20[d] | 0.53 | 0.170 | 1.11 | 0.224 | -0.26 | 0.733 |
| | | 0.41-0.60[c] | 1.99 | **1.E-06** | 0.62 | 0.489 | - | - |
| | | 0.61-0.80[c] | 1.14 | **0.018** | -0.43 | 0.620 | - | - |
| | | > 0.80[c] | -0.87 | 0.538 | -0.74 | 0.378 | - | - |
| MALE | Income | > 1MW | 0.77 | **0.001** | 0.76 | **0.002** | 0.76 | **0.001** |
| | Genomic ancestry[a] | | -0.02 | 0.750 | 0.03 | 0.585 | -0.20 | 0.317 |
| | Genomic ancestry[b] | | | | | | | |
| | | 0.21-0.40[c] \|> 0.20[d] | 0.46 | 0.195 | 0.44 | 0.635 | 0.47 | 0.488 |
| | | 0.41-0.60[c] | -0.08 | 0.819 | 0.43 | 0.644 | - | - |
| | | 0.61-0.80[c] | -0.08 | 0.855 | 0.30 | 0.740 | - | - |
| | | > 0.80[c] | -1.18 | 0.411 | 0.42 | 0.633 | - | - |

Multiple linear regression adjusted by sex, income, and ancestry ($\beta_0$ for African ancestry = 22.85, $\beta_0$ for European ancestry = 25.67, and $\beta_0$ for Native American ancestry = 22.71 kg/m2); [a]Ancestry value in tenths; [b]Ancestry in classes of 0.20 units of ancestry, where reference on regression analysis is the class of ancestry 0-0.20; [c]For African and European ancestries there are five intervals, [d]For Native American ancestry there are two intervals. Values in bold indicate p-value < 0.05. MW = Minimum wage.

**Table S4. Multiple linear regression analyses of BMI on African, European, and Native American ancestries for older adults from Bambui cohort (n=1342).**

| Variables | | African | | European | | Native American | |
|---|---|---|---|---|---|---|---|
| | | β | p-value | β | p-value | β | p-value |
| Age (years) | | -0.11 | **2.0e-10** | -0.11 | **2.0e-10** | -0.11 | **4.0e-10** |
| Sex | Male | -2.04 | **9.0e-16** | -2.04 | **9.0e-16** | -2.03 | **1.0e-15** |
| Educational level | | | | | | | |
| | >= High school | 1.12 | **2.0e-04** | 1.10 | **3.0e-04** | 1.09 | **4.0e-04** |
| Income | >= USD180 | 0.39 | 0.180 | 0.39 | 0.183 | 0.38 | 0.187 |
| Genomic ancestry[a] | | -0.06 | 0.587 | 0.06 | 0.501 | -0.23 | 0.548 |
| Interaction: ancestry*male | | -0.28 | 0.123 | 0.19 | 0.233 | 0.69 | 0.272 |
| Genomic ancestry[b] | | | | | | | |
| | 0.21-0.40[c] \| (> 0.20)[d] | -0.92 | **0.013** | 2.54 | 0.089 | -2.74 | **0.027** |
| | 0.41-0.60[c] | 0.81 | 0.329 | 2.58 | **0.011** | - | - |
| | 0.61-0.80[c] | 1.44 | 0.455 | 1.84 | **0.039** | - | - |
| | > 0.80[c] | -2.35 | **0.013** | 2.39 | **0.007** | - | - |

Multiple linear regression adjusted by age, sex, educational level, income, and ancestry ($\beta_0$ for African ancestry = 33.21, $\beta_0$ for European ancestry = 32.67, $\beta_0$ for Native American ancestry = 33.36 kg/m2); [a]Ancestry value in tenths; [b]Ancestry in classes of 0.20 units of ancestry, where reference on regression analysis is the class of ancestry 0-0.20; [c]For African and European ancestries there are five intervals, [d]For Native American ancestry there are two intervals. The value of USD 180 in Bambui corresponds to the median of income in Bambui cohort. Values in bold indicate p-value < 0.05.

**Table S5. Genome-wide p-values thresholds obtained for each vector of local ancestries (0, 1, or 2 chromosomes of given ancestry) for the entire dataset, obtained following the method of Shriner et al. (2011) based on the estimation of the effective number of tests (ENT). A.** Salvador, **B.** Pelotas, **C.** Bambuí. The number of tests was estimated using PCAdmix inferences.

**A. SALVADOR**

| | Ancestry | ENT | p-value[a] |
|---|---|---|---|
| FEMALE + MALE | African | 440.34 | 1.13e-04 |
| | European | 284.25 | 1.75e-04 |
| | Native American | 1038.72 | 4.82e-05 |

**B. PELOTAS**

| | Ancestry | ENT | p-value[a] |
|---|---|---|---|
| FEMALE + MALE | African | 1148.14 | 4.35e-05 |
| | European | 736.12 | 6.79e-05 |
| | Native American | 1124.93 | 4.44e-05 |
| FEMALE | African | 1129.55 | 4.42e-05 |
| | European | 713.67 | 7.00e-05 |
| | Native American | 1112.04 | 4.49e-05 |
| MALE | African | 1166.44 | 4.28e-05 |
| | European | 758.21 | 6.59e-05 |
| | Native American | 1137.61 | 4.39e-05 |

**C. BAMBUI**

| | Ancestry | ENT | p-value[a] |
|---|---|---|---|
| FEMALE + MALE | African | 980.94 | 5.09e-05 |
| | European | 693.37 | 7.27e-05 |
| | Native American | 1184.35 | 4.21e-05 |

[a] Bonferroni p-value threshold for significance obtained by dividing 0.05 by ENT

**Table S6. Parental allele frequencies of SNPs considered significant at fine mapping analysis of consensus significant peaks in young adults of Pelotas cohort.** We considered significant, associations with p-values less than the ones obtained for the admixture mapping peaks. Lead SNP in bold. All SNPs were imputed.

| SNP | Effect allele | Effect allele frequency | Europeans | Africans | | | | East Asians | Americans | |
|---|---|---|---|---|---|---|---|---|---|---|
| | | | EUR | GWD | WAFR | EAFR | AFR | EAS | LAT | NAT |
| **Peak at 16q12.1 (African ancestry)** | | | | | | | | | | |
| rs76416629 | G | 0.090 | 0.007 | 0.367 | 0.421 | 0.419 | 0.42 | 0.193 | 0.066 | 0.100 |
| **Peak at 13q12.3 (European ancestry, females)** | | | | | | | | | | |
| **rs114066381** | A | 0.008 | 0.004 | 0.020 | 0.032 | 0.000 | 0.017 | 0 | 0.003 | - |
| rs113214936 | G | 0.020 | 0.001 | 0.146 | 0.134 | 0.232 | 0.181 | 0 | 0.01 | 0 |
| **Peak at 20p12.1-2 (European ancestry, females)** | | | | | | | | | | |
| **rs138074843** | A | 0.005 | 0.001 | 0.013 | 0.005 | 0.020 | 0.012 | 0 | 0 | - |
| rs139525158 | C | 0.005 | 0.001 | 0 | 0 | 0 | 0 | 0 | 0.040 | 0.073 |
| rs79474139 | G | 0.013 | 0 | 0.097 | 0.139 | 0.081 | 0.111 | 0 | 0.007 | - |

Allele frequency from 1000 Genomes Public Database, EUR: European (CEU, FIN, GBR, TSI and IBS), GWD: Gambian in Western Divisions in the Gambia, WAFR: Western African (YRI), EAFR: Eastern African (LWK), AFR: African (LWK and YRI), EAS: East Asian (CDX, CHB, CHS, JPT and KHV), LAT: Admixed Latin Americans (PUR, CLM, MXL and PEL).

**Table S7. Fine mapping of consensus significant peaks in young adults of Pelotas cohort.** We considered significant, associations with p-values less than the ones obtained for the admixture mapping peaks. Lead SNP in bold. All SNPs were imputed.

| SNP | Position (bp)[a] | Functional consequence | Gene or closest gene | Effect allele | β | SE | p-value |
|---|---|---|---|---|---|---|---|
| **Peak at 16q12.1 (African ancestry)** | | | | | | | |
| rs76416629 | 50726760 | 2kb upstream | *NOD2* | G | -1.025 | 0.1864 | 4.09e-08 |
| **Peak at 13q12.3 (European ancestry, females)** | | | | | | | |
| **rs114066381** | 30632736 | intergenic | *LINC00544, LINC00365* | A | 3.99 | 0.8487 | 2.76e-06 |
| rs113214936 | 30018418 | intron | *MTUS2* | G | 2.48 | 0.5726 | 1.51e-05 |
| **Peak at 20p12.1-2 (European ancestry, females)** | | | | | | | |
| **rs138074843** | 15933795 | intron | *MACROD2,LOC613266* | A | 4.231 | 1.026 | 3.90e-05[b] |
| rs139525158 | 15385339 | intron | *MACROD2* | C | 4.527 | 1.104 | 4.30e-05[b] |
| rs79474139 | 15034071 | intron | *MACROD2* | G | 2.76 | 0.699 | 8.19e-05[b] |

[a]bp: base pair (hg19). [b]Suggestive association.

**Table S8. Parental allele frequencies of SNPs considered significant at fine mapping analysis of consensus significant peaks in children of Salvador cohort.** We considered significant, associations with p-values less than the ones obtained for the admixture mapping peaks. Lead SNP in bold. All SNPs were imputed.

| SNP | Effect allele | Effect allele frequency | Europeans | Africans | | | | East Asians | Americans | |
|---|---|---|---|---|---|---|---|---|---|---|
| | | | EUR | GWD | WAFR | EAFR | AFR | EAS | LAT | NAT |
| **Peak at 10q22.1 (African ancestry)** | | | | | | | | | | |
| rs1334909357 | CTTT | 0.221 | 0.082 | 0.398 | 0.417 | 0.379 | 0.399 | 0.320 | 0.208 | - |
| **Peak at 10q22.3 (European and African ancestries)** | | | | | | | | | | |
| **rs79947827** | A | 0.006 | 0 | 0 | 0.019 | 0.005 | 0.012 | 0 | 0.003 | 0 |
| rs141274185 | T | 0.007 | 0 | 0 | 0.023 | 0.005 | 0.015 | 0 | 0.003 | 0 |

Allele frequency from 1000 Genomes Public Database, EUR: European (CEU, FIN, GBR, TSI and IBS), GWD: Gambian in Western Divisions in the Gambia, WAFR: Western African (YRI), EAFR: Eastern African (LWK), AFR: African (LWK e YRI), EAS: East Asian (CDX, CHB, CHS, JPT and KHV), LAT: Admixed Latin Americans (PUR, CLM, MXL and PEL), NAT: Native Americans (Chopccas, Matses, Moche and Uros).

**Table S9. Fine mapping of consensus significant peaks in children of Salvador cohort.** We considered significant associations with p-values less than the ones obtained for the admixture mapping peaks. Lead SNP in bold. All SNPs were imputed.

| SNP | Position (bp) [a] | Functional Consequence | Gene or Closest gene | Effect allele | β | SE | p-value |
|---|---|---|---|---|---|---|---|
| **Peak at 10q22.1 (African ancestry)** | | | | | | | |
| rs1334909357 | 72224380 | intron (LCR[b]) | *PALD1* | CTTT | 0.4701 | 0.1009 | 3.59e-06 |
| **Peak at 10q22.3 (European and African ancestries)** | | | | | | | |
| **rs79947827** | 80998528 | intron | *ZMIZ1* | A | 2.099 | 0.5035 | 3.28e-05 |
| rs141274185 | 80993437 | intron | *ZMIZ1* | T | 1.808 | 0.4516 | 6.59e-05 |

[a]bp: base pair (hg19). [b]Designated as LCR (Low Complexity Region) in GnomAD browser (https://gnomad.broadinstitute.org/variant/10-72224380-CTTT-C).

**Table S10. Classification of all females and of rs114066381-A carrier females by BMI categories.** BMI was classified according to the WHO guidelines: Underweight (BMI<18.5), Normal weight (>=18.5 and <25), Overweight (>=25 and <30), Obese (>=30 and <=40), and Morbidly obese (>40).

| | SAMPLE | N alleles | Low-weight | Normal | Overweight | Obese | Morbidly obese |
|---|---|---|---|---|---|---|---|
| PELOTAS | All Unrelated Females | 3590 | 0.07 (258) | 0.65 (2344) | 0.18 (660) | 0.08 (294) | 0.01 (38) |
| | *rs114066381-A carriers / All Unrelated Females* | *31/3590 (0.008)* | *2/258 (0.007)* | *11/2344 (0.004)* | *7/660 (0.010)* | *7/294 (0.023)* | *4/38 (0.10)* |
| SÃO PAULO | All Unrelated Females | 1302 | 0.02 (26) | 0.23 (298) | 0.37 (480) | 0.35 (462) | 0.03 (36) |
| | *rs114066381-A carriers / All Unrelated Females* | *25/1302 (0.019)* | *0 (0)* | *4/298 (0.013)* | *9/480 (0.018)* | *9/462 (0.019)* | *3/36 (0.083)** |
| BAMBUI | All Unrelated Females | 1032 | 0.07 (70) | 0.38 (394) | 0.38 (390) | 0.16 (166) | 0.01 (12) |
| | *rs114066381-A carriers / All Unrelated Females* | *11/1032 (0.010)* | *0 (0)* | *5/394 (0.012)* | *3/390 (0.007)* | *2/166 (0.012)* | *1/12 (0.083)** |
| BAMBUI | All Females | 1642 | 0.07 (108) | 0.38 (630) | 0.38 (620) | 0.16 (262) | 0.01 (22) |
| | *rs114066381-A carriers / All Females* | *16/1642 (0.009)* | *0 (0)* | *8/630 (0.012)* | *4/620 (0.006)* | *2/262 (0.007)* | *2/22 (0.09)** |
| BAMBUI + SÃO PAULO | All Unrelated Females | 2334 | 0.04 (96) | 0.30 (692) | 0.37 (870) | 0.27 (628) | 0.02 (48) |
| | *rs114066381-A carriers / All Unrelated Females* | *36/2334 (0.015)* | *0 (0)* | *9/692 (0.013)* | *12/870 (0.013)* | *11/628 (0.017)* | *4/48 (0.083)* |
| PUERTO RICO | All Unrelated Females | 2092 | 0.01 (26) | 0.32 (674) | 0.39 (804) | 0.26 (538) | 0.02 (50) |
| | *rs114066381-A carriers / All Unrelated Females* | 12/2092 *(0.005)* | *0 (0)* | *4 (0.006)* | *1 (0.001)* | *7 (0.013)* | *0 (0)* |
| SOWETO | All Unrelated Females | 1832 | 0.10 (184) | 0.45 (832) | 0.27 (486) | 0.17 (306) | 0.01 (24) |
| | *rs114066381-A carriers / All Unrelated Females* | *31/1832 (0.017)* | *1/184 (0.005)* | *9/832 (0.011)* | *12/486 (0.024)* | *8/306 (0.026)* | *1/24 (0.041)* |

*The woman with the highest BMI in the cohort has the rs114066381-A allele.

**Table S11. Association of GWAS-catalog-Hits on three Brazilian cohorts.** Results of replication analysis showing 28 associated with BMI in Pelotas and Bambui cohorts

| CHR | SNPS | Allele | Gene | GWAS-catalog | | | Pelotas | | | Bambui | | |
|---|---|---|---|---|---|---|---|---|---|---|---|---|
| | | | | Freq | ME | Age | Freq | ME(SE) | p-BHa | Freq | ME(SE) | p-BHa |
| 1 | rs3843262 | A/T | intergenic | 0.43 | 0.03 | Adult | 0.48 | 0.15(0.1) | 4.03e-01 | 0.45 | 0.63(0.18) | **3.58e-02 (A,F)** |
| 1 | rs12401738 | A/G | GIPC2,DNAJB4 | 0.34 | 0.02 | Adult | 0.25 | 0.23(0.12) | 2.50e-01 | 0.23 | -0.88(0.2) | **2.67e-03 (A,F)** |
| 1 | rs4130548 | C/T | GIPC2,DNAJB4 | 0.35 | 0.03 | Adult | 0.25 | 0.24(0.12) | 2.32e-01 | 0.23 | -0.84(0.2) | **3.45e-03 (A,F)** |
| 3 | rs13078960 | T/G | CADM2 | 0.81 | 0.03 | Adult | 0.83 | -0.09(0.14) | **2.33e-02 (M)** | 0.79 | -0.69(0.26) | 1.04e-01 |
| 3 | rs9852127 | A/G | CADM2 | 0.19 | 0.03 | Adult | 0.17 | 0.09(0.14) | **2.33e-02 (M)** | 0.22 | 0.63(0.26) | 1.52e-01 |
| 3 | rs9852859 | C/T | CADM2 | 0.19 | 0.03 | Adult | 0.17 | 0.09(0.14) | **2.21e-02 (M)** | 0.21 | 0.69(0.26) | 1.04e-01 |
| 3 | rs9818122 | C/T | CADM2,CADM2-AS2 | 0.20 | 0.03 | Adult | 0.20 | 0.05(0.13) | **1.50e-02 (M)** | 0.25 | 0.48(0.24) | 2.88e-01 |
| 3 | rs13078807 | G/A | CADM2 | 0.20 | 0.10 | Adult,Children | 0.16 | 0.08(0.14) | **3.71e-02 (M)** | 0.21 | 0.69(0.26) | 1.04e-01 |
| 11 | rs7113874 | C/T | TRIM66 | 0.64 | 0.03 | Adult | 0.58 | 0.03(0.1) | 4.01e-01 | 0.61 | 0.33(0.18) | **3.78e-02 (F)** |
| 12 | rs7138803 | A/G | intergenic | 0.38 | 0.12 | Adult,Children | 0.29 | 0.36(0.11) | **2.52e-02 (A,M)** | 0.31 | 0.3(0.21) | 4.79e-01 |
| 16 | rs12444979 | C/T | intergenic | 0.87 | 0.17 | Adult | 0.89 | 0.51(0.16) | **2.52e-02 (A)** | 0.90 | -0.8(0.33) | 1.61e-01 |
| 16 | rs12446632 | G/A | intergenic | 0.87 | 0.05 | Adult,Children | 0.89 | 0.51(0.16) | **2.52e-02 (A)** | 0.90 | -0.77(0.33) | 1.75e-01 |
| 16 | rs9930333 | G/T | FTO | 0.43 | 0.08 | Adult | 0.44 | 0.53(0.1) | **1.32e-05(A,M)** | 0.44 | 0.44(0.19) | 1.75e-01 |
| 16 | rs1558902 | A/T | FTO | 0.42 | 0.39 | Adult,Children | 0.36 | 0.73(0.11) | **3.26e-09 (A,F,M)** | 0.38 | 0.45(0.19) | 1.75e-01 |
| 16 | rs62033400 | G/A | FTO | - | 0.08 | Adult | 0.35 | 0.66(0.11) | **7.49e-08 (A,M)** | 0.36 | 0.53(0.2) | 1.04e-01 |
| 16 | rs8050136 | A/C | FTO | 0.41 | 0.09 | Adult,Children | 0.41 | 0.54(0.1) | **1.08e-05 (A,M)** | 0.41 | 0.47(0.19) | 1.52e-01 |
| 16 | rs3751812 | T/G | FTO | 0.40 | 0.08 | Adult | 0.34 | 0.71(0.11) | **9.28e-09 (A,F,M)** | 0.36 | 0.49(0.2) | 1.52e-01 |
| 16 | rs9939609 | A/T | FTO | 0.41;0.13 | 0.33 | Adult,Children | 0.42 | 0.52(0.1) | **2.45e-05 (A,M)** | 0.41 | 0.42(0.19) | 1.96e-01 |
| 17 | rs16943469 | T/A | YPEL2 | 0.02 | 3.98 | Adult | 0.99 | -0.12(0.43) | 6.84e-01 | 0.99 | -4.87(1.61) | **2.89e-03 (F)** |
| 18 | rs6567160 | C/T | intergenic | 0.24 | 0.09 | Adult,Children | 0.20 | 0.58(0.13) | **2.25e-04 (A,M)** | 0.18 | 0.32(0.25) | 3.68e-01 |
| 18 | rs571312 | A/C | intergenic | 0.24 | 0.23 | Adult,Children | 0.22 | 0.49(0.12) | **1.83e-04 (A,M)** | 0.20 | 0.16(0.24) | 7.51e-01 |
| 18 | rs17782313 | C/T | intergenic | 0.21 | 0.20 | Adult | 0.22 | 0.46(0.13) | **6.32e-04 (A,M)** | 0.20 | 0.09(0.25) | 7.51e-01 |
| 18 | rs12969709 | A/C | intergenic | 0.26 | 0.06 | Adult | 0.21 | 0.52(0.13) | **5.49e-04 (A,M)** | 0.19 | 0.3(0.24) | 4.07e-01 |
| 18 | rs11663816 | C/T | intergenic | 0.27 | 0.06 | Adult | 0.20 | 0.53(0.13) | **9.05e-04 (A,M)** | 0.18 | 0.29(0.25) | 4.15e-01 |
| 19 | rs757318 | C/A | CRTC1 | 0.51 | 0.02 | Adult | 0.46 | 0.24(0.1) | **1.38e-03 (M)** | 0.44 | -0.06(0.18) | 9.18e-01 |
| 19 | rs11672660 | C/T | GIPR | 0.79 | 0.04 | Adult | 0.84 | 0.1(0.14) | 5.11e-01 | 0.85 | 0.84(0.26) | **3.58e-02 (A,F)** |
| 19 | rs1800437 | G/C | GIPR | 0.79 | 0.03 | Adult | 0.84 | 0.1(0.14) | 5.04e-01 | 0.85 | 0.85(0.26) | **3.58e-02 (A,F)** |
| 19 | rs2287019 | C/T | QPCTL | 0.80 | 0.15 | Adult | 0.85 | 0.13(0.14) | 4.96e-01 | 0.86 | 0.87(0.26) | **3.58e-02 (A,F)** |

CHR: chromosome; RA: risk allele; ME: Main effect (kg/m2); SE: Standard error of ME; Freq: Allele frequency; p-BH: Benjamini Hochberg adjusted p-value. aP-value statistically significant stratified by sex: A: all individuals, F: females, and M: males.. BH p-values in bold are < 0.05

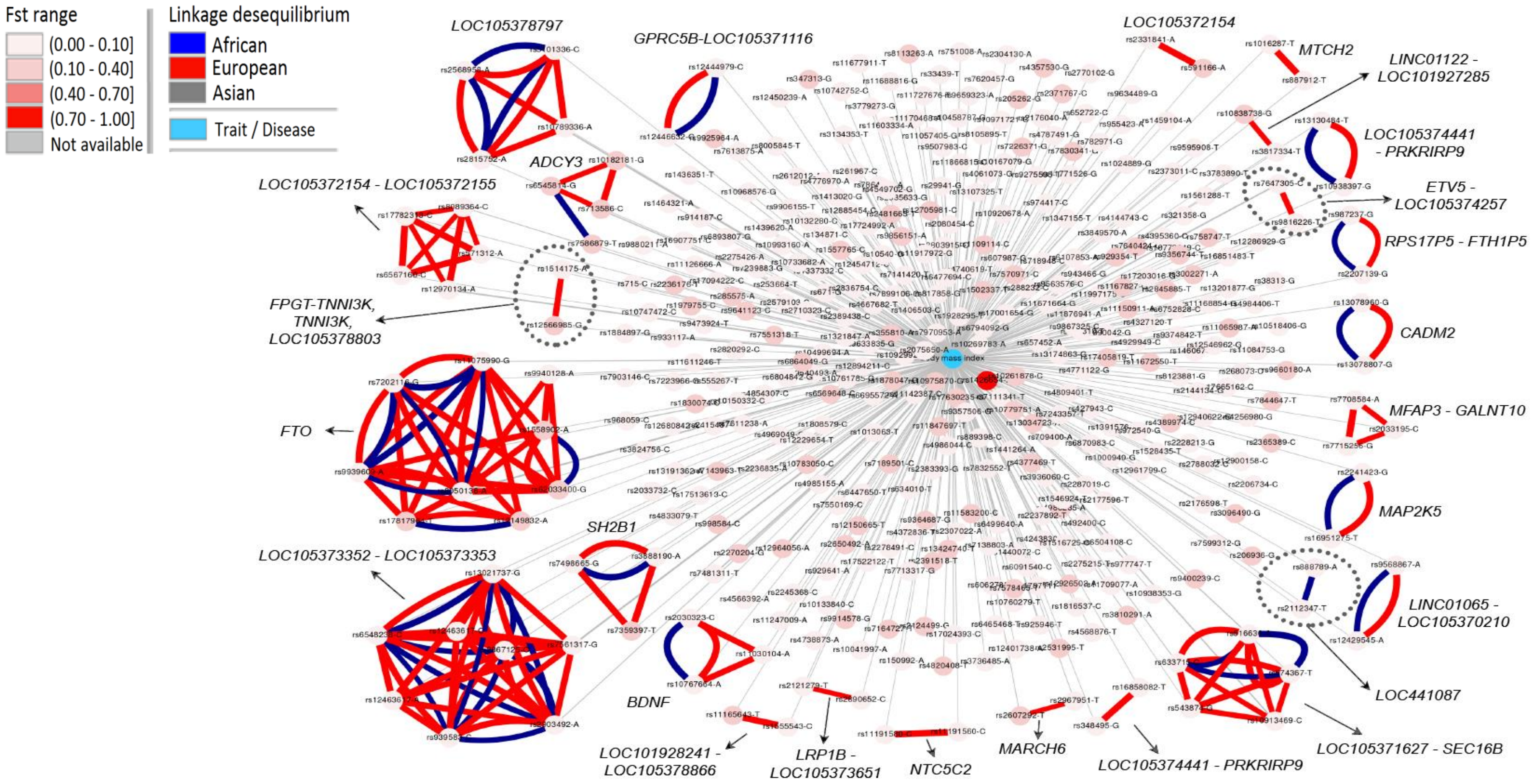


Figure S1. Allelic architecture of Body Mass Index. SNP-Disease Network for Body Mass Index (BMI) considering African and European populations. BMI is represented as a blue node and GWAS hits (SNPs) as red nodes. Red color gradient of the SNP nodes denotes the level of differentiation (FST) of the allele frequency of the risk allele between European and Africans, based on 1000 Genomes Project. Among the 389 SNPs associated with BMI, we observed 24 clusters of risk SNPs that show high linkage disequilibrium (r2 >= 0.7) between Africans (blue edges) and Europeans (red edges). The SNP-Disease Network was obtained from the DANCE web tool (www.ldgh.com.br/dance). We edited the figure to include arrows and gene names. Filters used in DANCE to reproduce the visualization: Network = SNP-Disease Network, Ancestry = FST - African x European, Phenotype = body mass index and linkage disequilibrium in Africans and Europeans.

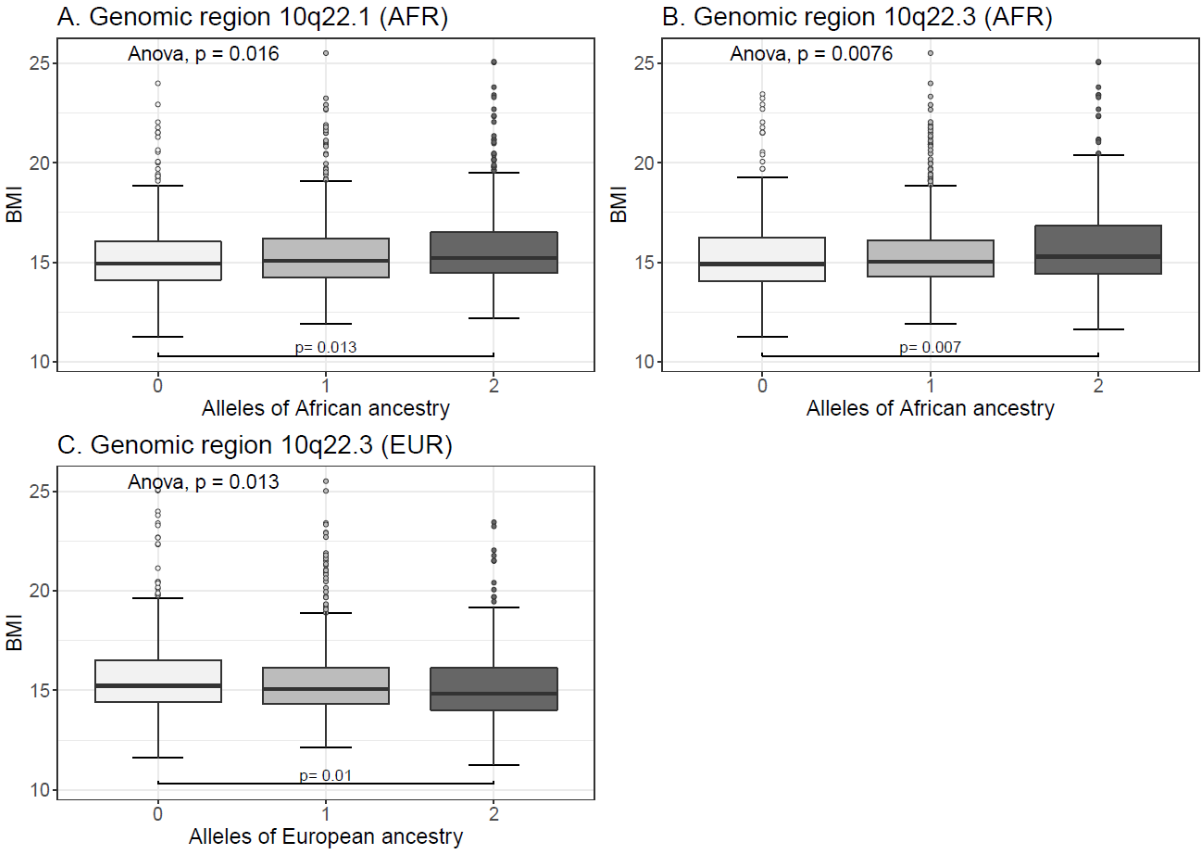


**Figure S2. Distribution of BMI in the children cohort of Salvador for each allele of African ancestry at A. 10q22.1 and B. 10q22.3, and of European ancestry at C. 10q22.3**. Plots for the lead window (i.e. genomic window with the lowest p-value in the linear regression result) of each genomic region. BMI mean values of the three different groups (0 alleles, 1 allele and 2 alleles) were compared using one-way analysis of variance (ANOVA). P-values obtained with these tests are depicted in each boxplot. Post-hoc test was performed using Tukey´s HSD test, statistically significant difference were observed only between the groups with 0 and 2 alleles of European ancestry on the respective genomic region.

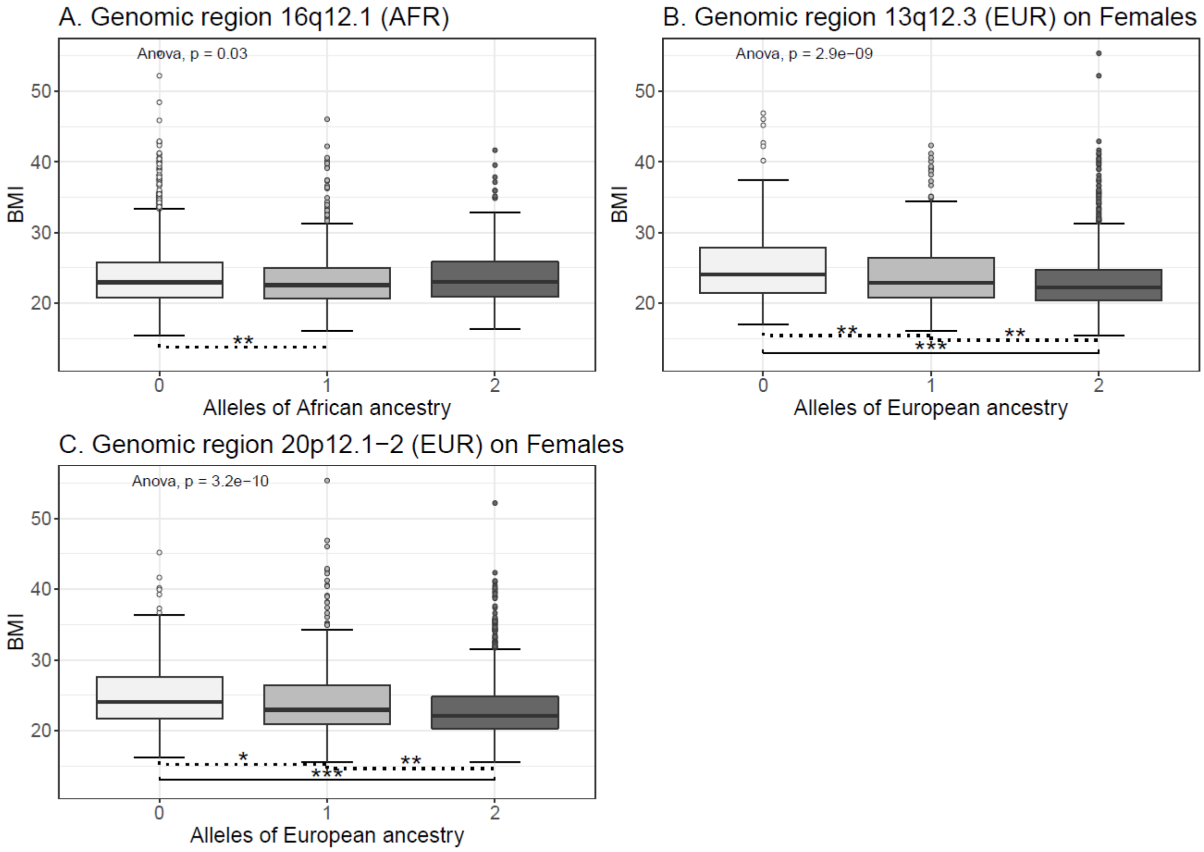


**Figure S3. Distribution of BMI in young adults from Pelotas cohort for each allele of African ancestry at A. 16q12.1, and European ancestry at B. 13q12.3 and C. 20p12.1-2.** Plots for the lead window (the genomic window with the lowest p-value in the linear regression result) of each genomic region. BMI mean values of the three different groups (0, 1 or 2 alleles) were compared using one-way analysis of variance (ANOVA). P-values obtained with these tests are depicted in each boxplot. Post-hoc test was performed using Tukey´s HSD test, p-values were represented as: *0.01 -0.05, ** <0.01, *** <0.001.

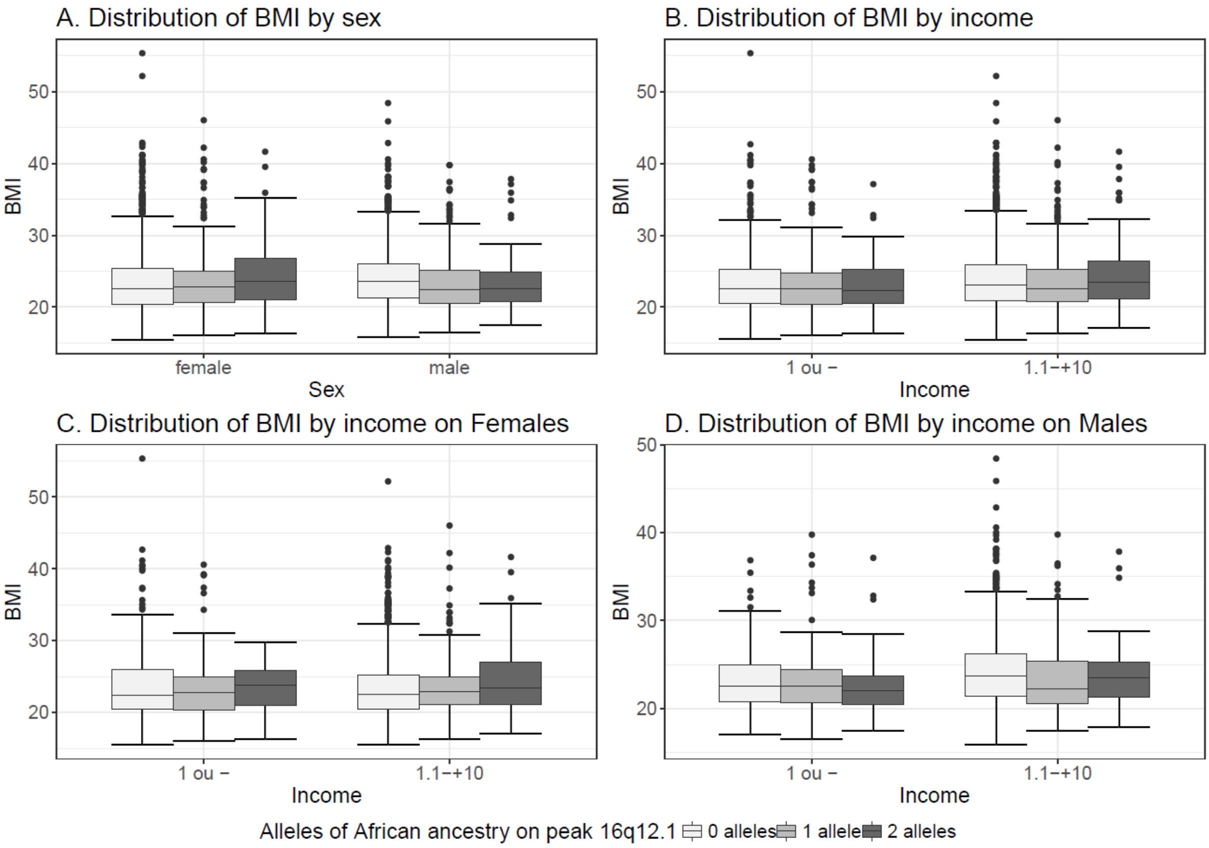


**Figure S4. Distribution of BMI for each allele of African ancestry 16q12.1 in young adults from Pelotas cohort stratified by A. sex, B. income, C. income on females, and D. income on males.**

**A.**Children from Salvador – African ancestry at 10q22.1 peak

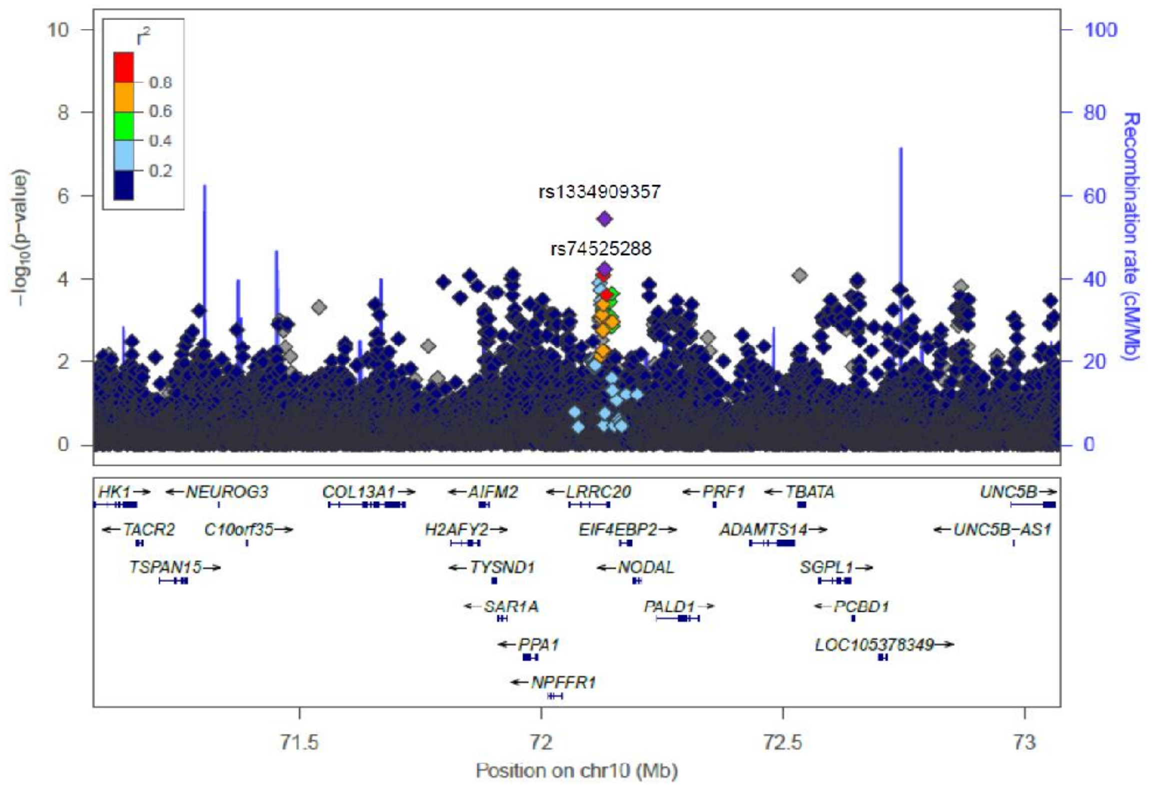


**B.** Children from Salvador – African and European ancestry at 10q22.3 peak

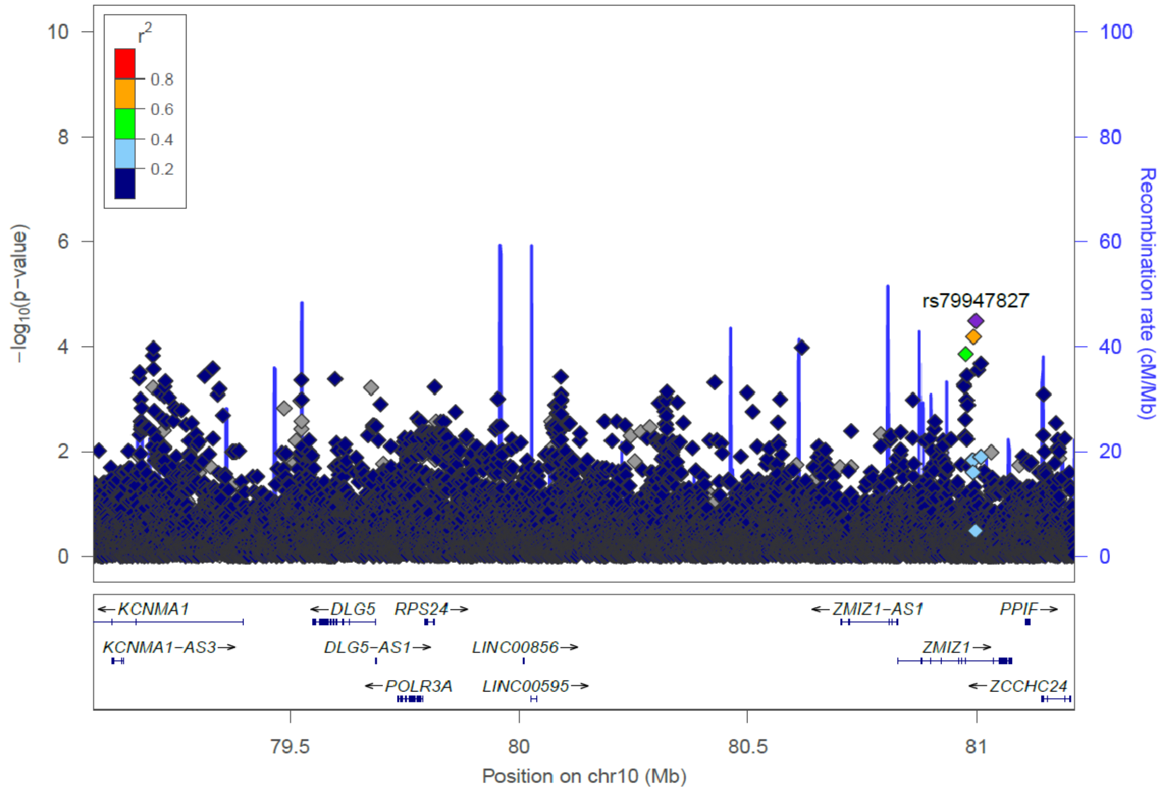

**C.** Young adults from Pelotas – African ancestry at 16q12.1 peak

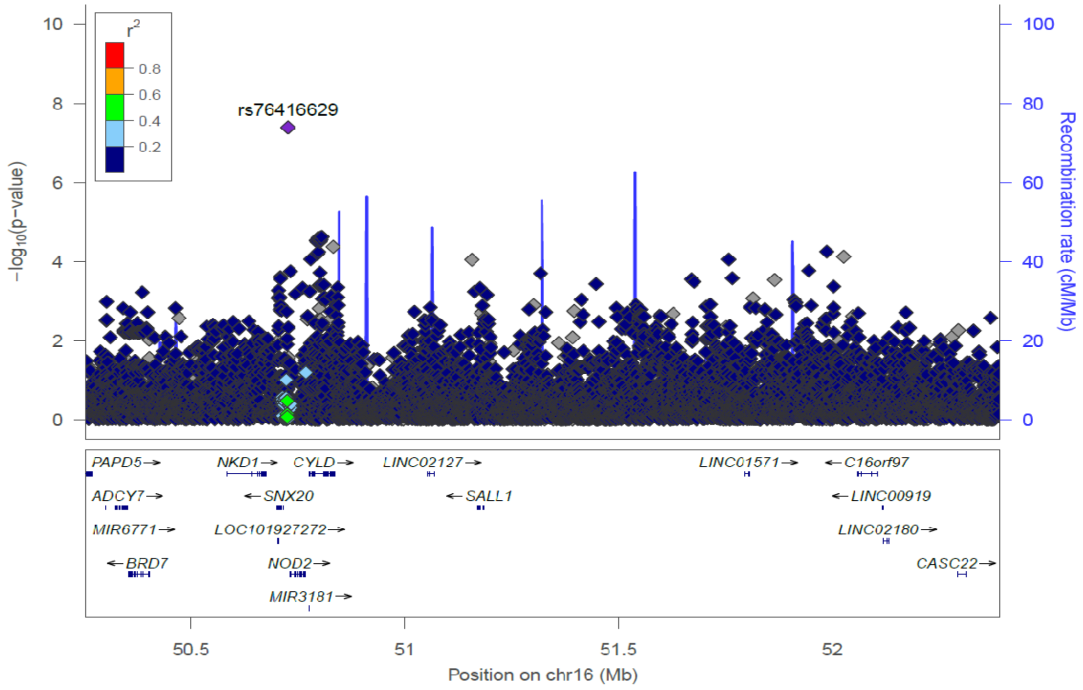


**D**. Young adults from Pelotas - European ancestry in females at 20p12.1-2 peak

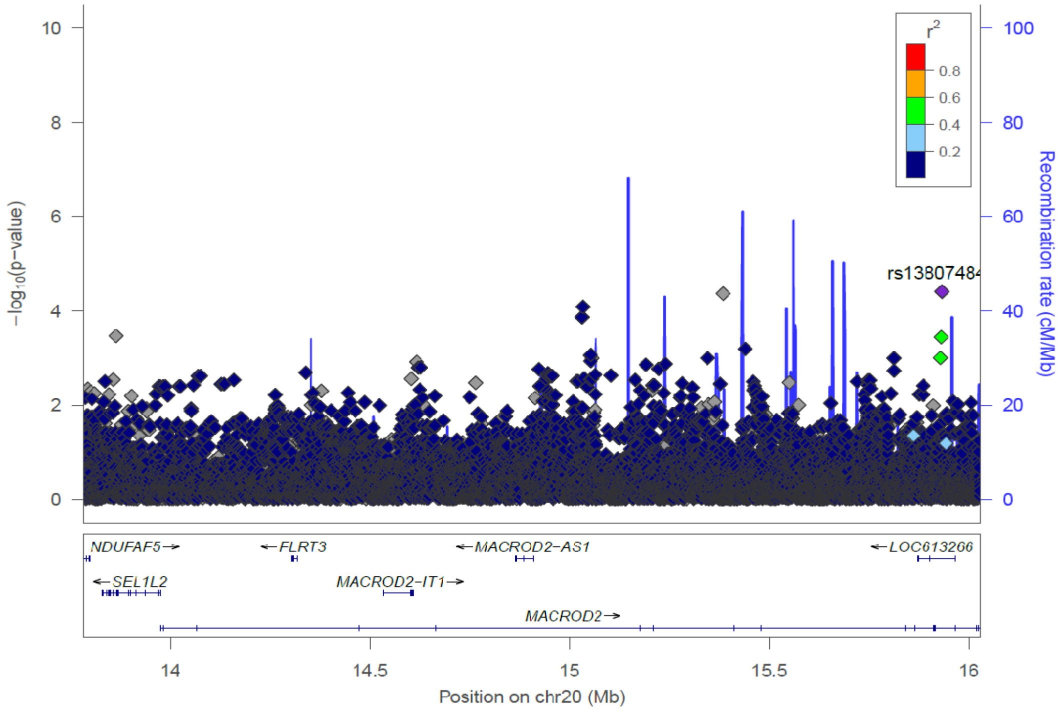


**Figure S5. LocusZoom plot of the fine-mapping of consensus significant Admixture Mapping peaks performed using both genotyped and imputed SNPs ±1 Mb from target region (lead windows).** The SNP with the lowest p-value is color coded in purple and labeled. The linkage disequilibrium between this SNP and the remaining nearby SNPs is indicated by the color coding according to $r^2$ values based on Africans from 1000 Genomes Project. **A**. Children from Salvador at Admixture peak 10q22.1 associated with African ancestry; **B.** Children from Salvador at admixture peak 10q22.3 associated with African and European ancestries; **C.** Young adults from Pelotas at Admixture peak 16q12.1 associated with African ancestry; **D.** Young adults from Pelotas at Admixture peak 20p12.1-2 associated with European ancestry in females only.

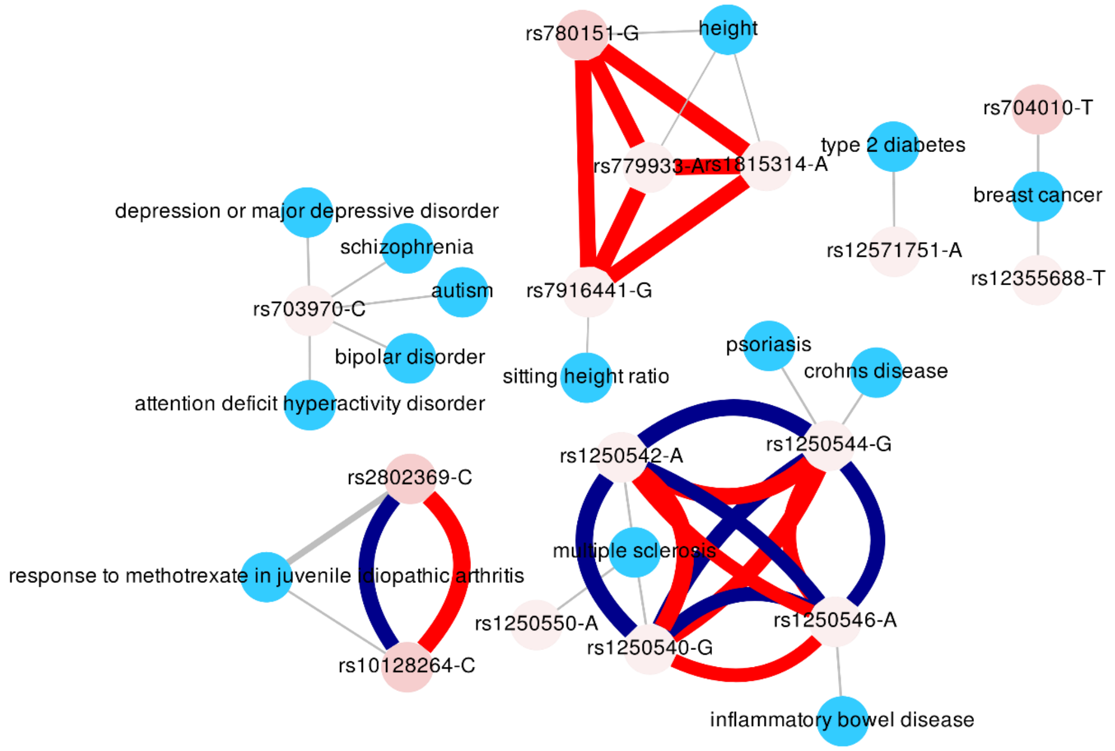


**Figure S6. DANCE Analysis.** SNP-Disease Network for ZMIZ1 gene considering African and European ancestries. ZMIZ1 is one of the genes most associated with complex phenotypes. DANCE analysis (Araújo et al., 2016) shows that SNPs in ZMIZ1 are associated with 14 complex disorders. On the network, complex phenotypes are represented as blue nodes and the genetic diversity ($F_{ST}$) between African and European is represented in the red gradient for each risk-allele. Also, the thickness of the edges represents linkage disequilibrium ($r^2$ >=0.7) for African (in dark blue) and Europeans (in red).

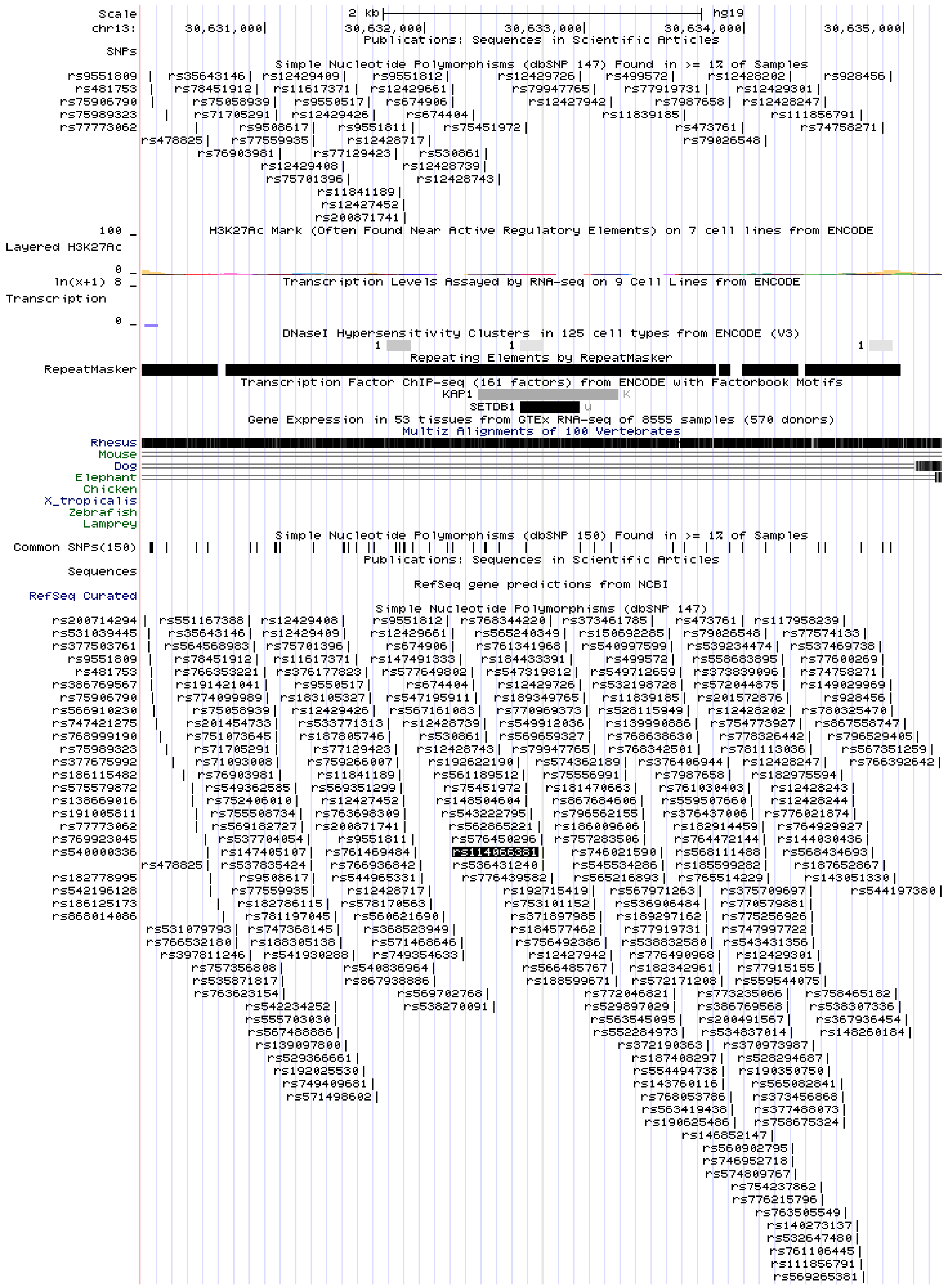

**Figure S7. Genomic context of rs114066381 by UCSC Genome Browser.** The SNP rs114066381 is located within a CTCF-binding site and contains binding sites for KAP1 and SETDB1 proteins, but none evidence of enhancer function is demonstrated by RNA-seq assays.

### Alignment block for hg38 chr13:30058574– 30058628 (54 bps)

(the full                                                     rs114066381

```
                   Human   CCTTTTA---------------
AAAAAAGCTATTTGCAGCGTTTATCAATTGGATAAAGTACATTTCCAT
                   Chimp   CCTTTTA---------------
AAAAAAGCTATTTGCAGCGTTTATCAATTGGATAAAGTACATTTCCAT
                  Bonobo   CCTTTTA---------------
AAAAAAGCTATTTGCAGCGTTTATCAATTGGATAAAGTACATTTCCAT
                 Gorilla   CCTTTTA---------------AAAAAAGCTATTTGCAG-
GTTTATCAATTGGATAAAGTACATTTCCAT
               Orangutan   CCTTTTT---------------
AAAAAAGCTATTTGCAGCTTTTAACAATTGGATAAAGTATATTTCCAT
                  Gibbon   CCTTTTT---------------
AAAAAAGCTATTTGCAGCTTTTAACAATTGGATAAAGTATATTTCCAT
                  Rhesus   CCTTTTT---------------
TAAAAAGCTATTTGCAGCTTTTAACAATTGGGTAAAGTACATTTCCAC
     Crab-eating macaque   CCTTTTT---------------
TAAAAAGCTATTTGCAGCTTTTAACAATTGGGTAAAGTACATTTCCAC
      Pig-tailed macaque   CCTTTTT---------------
TAAAAAGCTATTTGCAGCTTTTAACAATTGGGTAAAGTACATTTCCAT
            Green monkey   CCTTTTT---------------
TAAAAAGCTATTTGCAGCTTTTAACAATTGGGTAAAGTACATTTCCAT
         Angolan colobus   CCTTTTT---------------
AAAAAAGCTATTTGCAGCTTTTAACAATTGGGTAAAGTATATTTCCAT
Golden snub-nosed monkey   CCTTTTT---------------
AAAAAAGCTATTTGCAGCTTTTAACAATTGGGTAAAGTACATTTCCAT
 Black snub-nosed monkey   CCTTTTT----------------
TAAAAGCTATTTGCAGCTTTTAACAATTGGGTAAAGTACATTTCCAT
                Marmoset   CCTTTTTTTTTTT--
TTTAAAAAAAGAAACCATTTGAAGCTTTTAGTGATTGGGTAAAATACATTCCCAT
          Squirrel monkey   CCTTTTTTT----------
AAAAAAAAATATATTTGAAACTTTTAGCAATTGGGTAAAATACATTCCCAT
     White-faced sapajou   CCTTTTTTTTTT---------------
AACTATTTGAAGCTTTTCGCGATTGGGTAAAATACATTCCCAT
       Ma's night monkey
CCTTTTTATTTTTTCTTTTTTTAAAAAAACTATTTGAAGCTTTTAGTGATTGGGTAAAATACATTCCCAT
       Coquerel's sifaka
======================================================================
             Mouse lemur
======================================================================
          Sclater's lemur
======================================================================
             Black lemur
======================================================================
                   Mouse
======================================================================
               Armadillo
======================================================================
                     Dog
======================================================================
                Bushbaby
======================================================================
                  Baboon
======================================================================
                 Tarsier
======================================================================
                   Drill   ---------------------------------------------------------------------
-
           Sooty mangabey
======================================================================
        Proboscis monkey
======================================================================
```

**Figure S8. Multiple sequence alignment for the conserved block spanning rs114066381 in 30 mammal genomes.** A 54 base pairs region is shown for visualization purposes, the alignment blocks are available at https://genome.ucsc.edu/cgi-bin/hgc?hgsid=667750385_0jgbB4nkoSqTYMmifWlMnJ79EIRR&c=chr13&l=30057562&r=30059617&o=30058542&t=30058655&g=multiz30way&i=multiz30way.

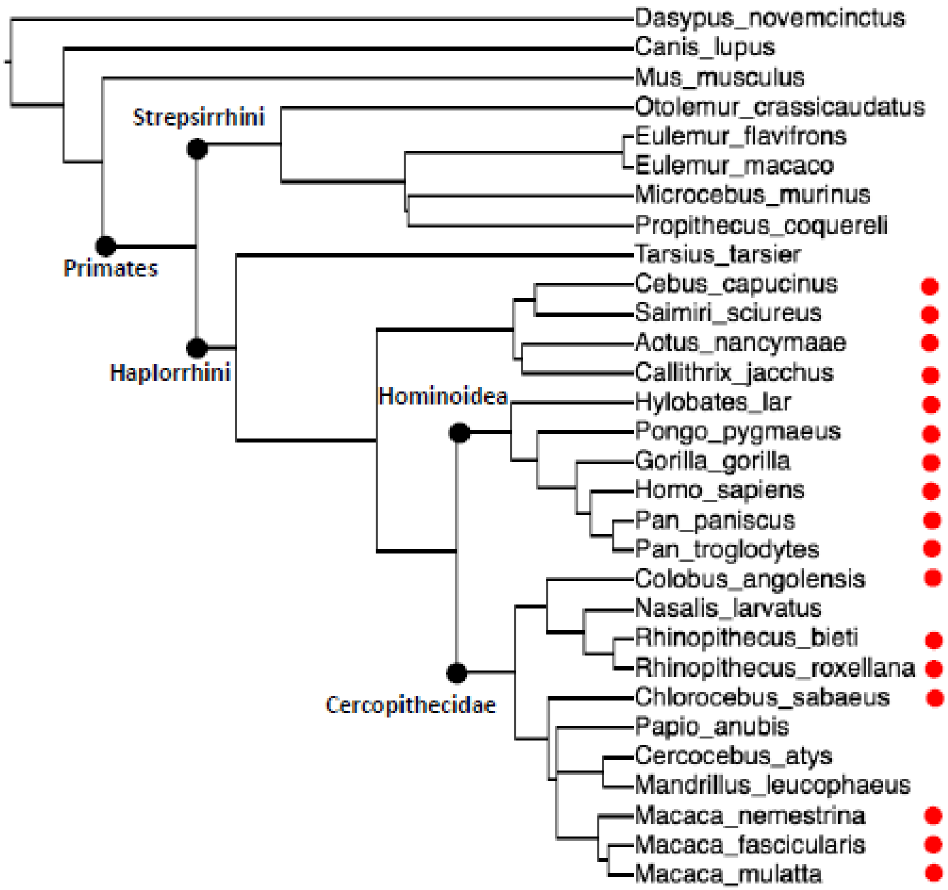


**Figure S9**. **Phylogenetic tree for the 30 mammal genomes evaluated for the presence of the homologous region spanning rs114066381.** Red dots indicate presence of conserved block spanning rs114066381 in species' genomes.

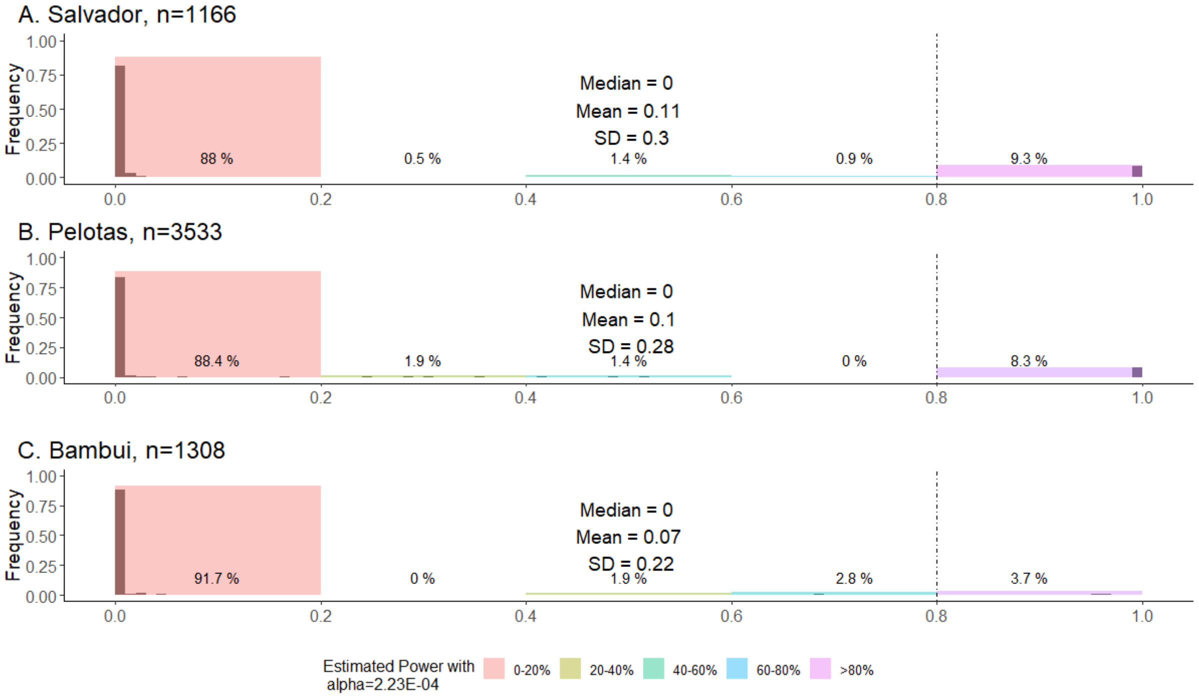


**Figure S10. Distribution of 216 GWAS-Hits power estimations on three Epigen Cohorts. Power was estimated for each individual SNP on each cohort according to the specific BMI distribution (mean ± SD).** Sample sizes correspond to individuals with no missing genotyped or imputed SNPs. Expected power for all 216SNPs in each population is represented by mean and median on each cohort. Dashed line represents 80% cut-off of statistical power. We used the software QUANTO v1.2.4 for all analysis with a α (alpha) value of 0.00023, as the probability of rejecting the true null hypothesis.

**A.**

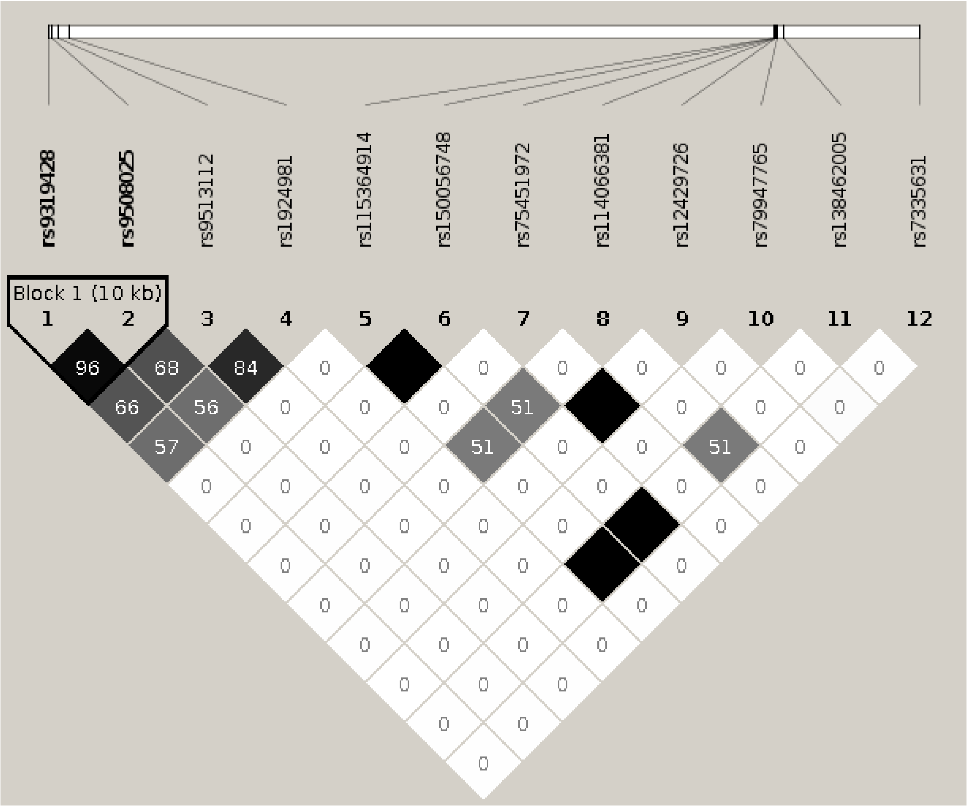


**B.**

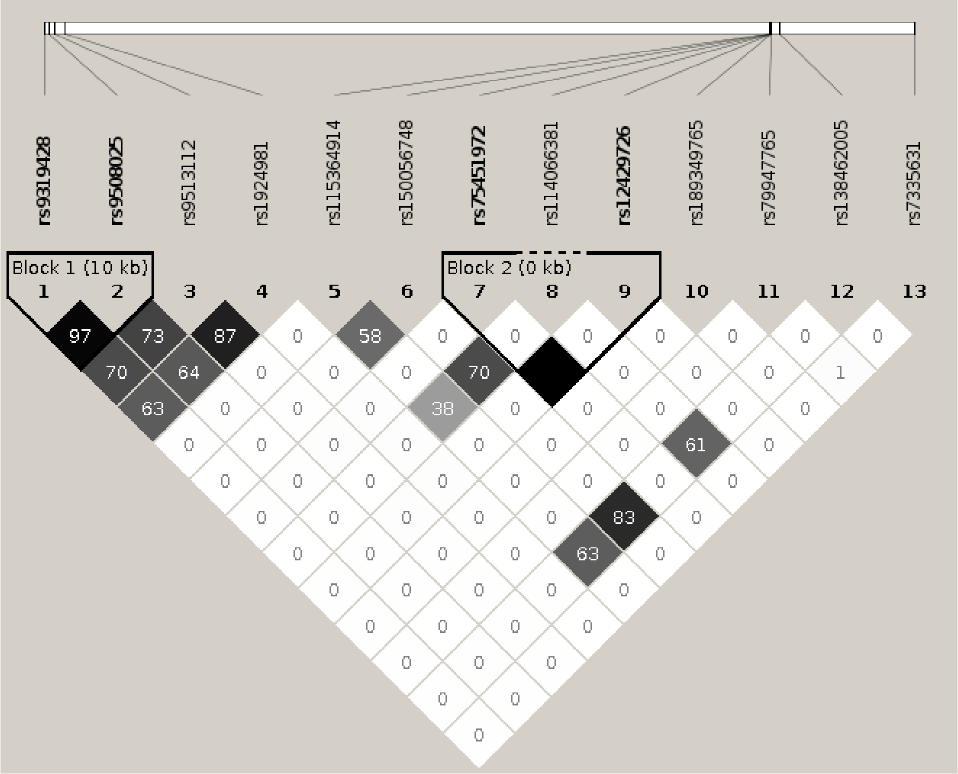


**Figure S11.** rs114066381 does not show LD ($r^2<0.7$) with any SNP mapped in close regulatory regions (in sensu HaploREG v4.1 in Ward and Kellis 2016) in **A.** Pelotas and in **B**. Bambui by Haploview.

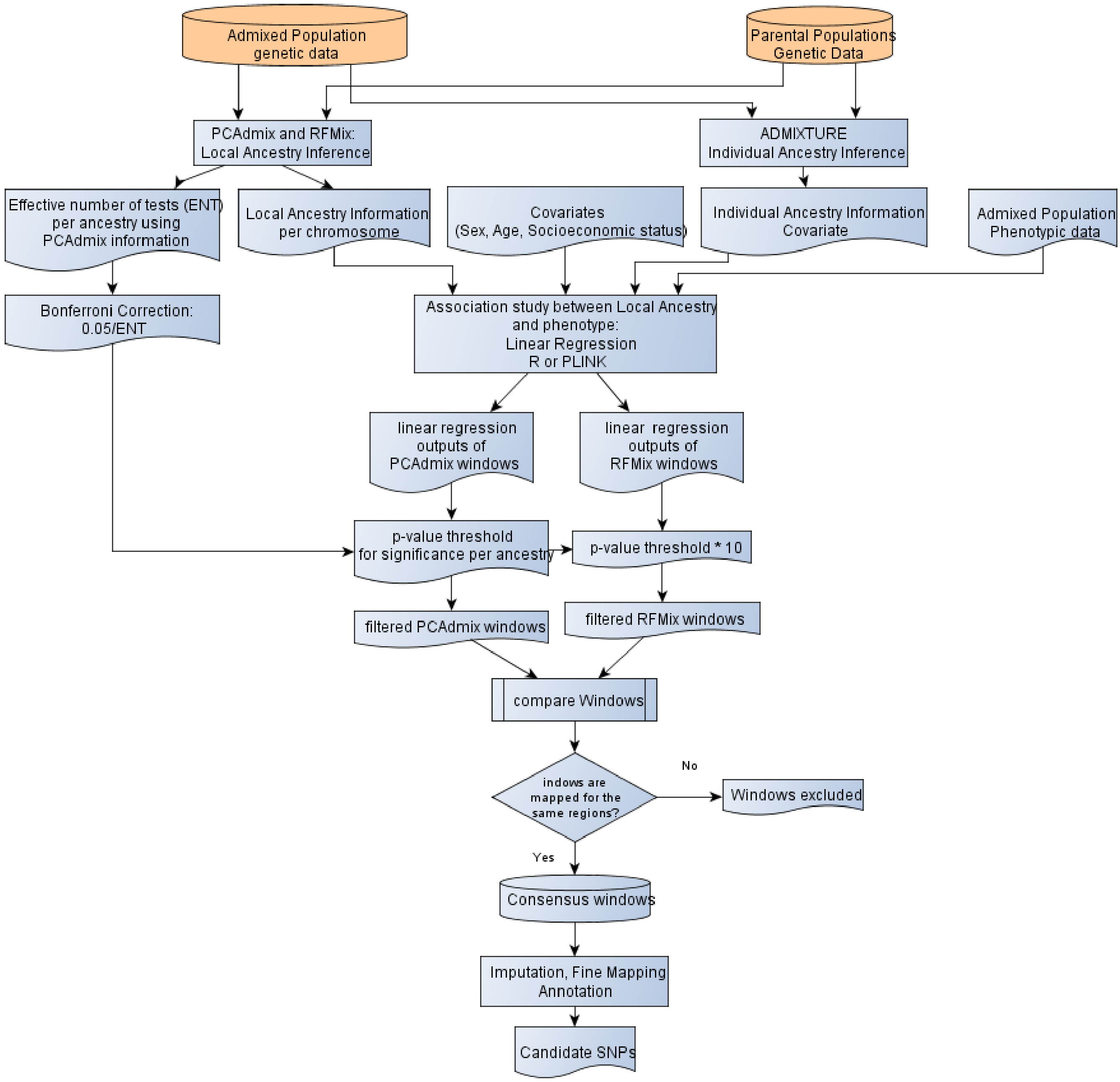


**Figure S12. Flowchart of the study design. Data requirements for admixture mapping include genotype and phenotype data for samples from the admixed population and genotype data from ancestral populations.** First, the individual and local ancestry of admixed population are inferred using ADMIXTURE, PCAdmix and RFMix (only for the chromosomes we found hits using PCAdmix) softwares. Then, an association of local ancestry with the phenotype is tested using a multiple regression including the individual ancestry and other descriptive information as covariates. Threshold p-value accounting for multiple testing was defined following the method of Shriner et al. (2011), based on the estimation of the effective number of tests (ENT) for each chromosome for each individual. This was used to filter the associated peaks of ancestry. In our approach, we compared the significant admixture mapping peaks of each chromosome obtained using PCAdmix with the significant admixture mapping peaks obtained using RFMix inferences for the same chromosomes, and, considered a consensus significant peak the ones mapped for the same chromosome region using both inferences. Only these consensus significant admixture mapping peaks were taken for further analyses of imputation, fine mapping and SNP annotation, generating a list of SNPs or genes associated with the phenotype.
